# Shotgun Transcriptome and Isothermal Profiling of SARS-CoV-2 Infection Reveals Unique Host Responses, Viral Diversification, and Drug Interactions

**DOI:** 10.1101/2020.04.20.048066

**Authors:** Daniel J. Butler, Christopher Mozsary, Cem Meydan, David Danko, Jonathan Foox, Joel Rosiene, Alon Shaiber, Ebrahim Afshinnekoo, Matthew MacKay, Fritz J. Sedlazeck, Nikolay A. Ivanov, Maria Sierra, Diana Pohle, Michael Zietz, Undina Gisladottir, Vijendra Ramlall, Craig D. Westover, Krista Ryon, Benjamin Young, Chandrima Bhattacharya, Phyllis Ruggiero, Bradley W. Langhorst, Nathan Tanner, Justyna Gawrys, Dmitry Meleshko, Dong Xu, Peter A. D. Steel, Amos J. Shemesh, Jenny Xiang, Jean Thierry-Mieg, Danielle Thierry-Mieg, Robert E. Schwartz, Angelika Iftner, Daniela Bezdan, John Sipley, Lin Cong, Arryn Craney, Priya Velu, Ari M. Melnick, Iman Hajirasouliha, Stacy M. Horner, Thomas Iftner, Mirella Salvatore, Massimo Loda, Lars F. Westblade, Melissa Cushing, Shawn Levy, Shixiu Wu, Nicholas Tatonetti, Marcin Imielinski, Hanna Rennert, Christopher E. Mason

**Author notes:** These authors contributed equally to this work (Daniel J. Butler, Christopher Mozsary, Cem Meydan, David Danko, Jonathan Foox). To whom correspondence should be addressed, **Corresponding Authors:** Marcin Imielinski; Hanna Rennert; Christopher E. Mason.

## Abstract

The Severe Acute Respiratory Syndrome Coronavirus 2 (SARS-CoV-2) has caused thousands of deaths worldwide, including >18,000 in New York City (NYC) alone. The sudden emergence of this pandemic has highlighted a pressing clinical need for rapid, scalable diagnostics that can detect infection, interrogate strain evolution, and identify novel patient biomarkers. To address these challenges, we designed a fast (30-minute) colorimetric test (LAMP) for SARS-CoV-2 infection from naso/oropharyngeal swabs, plus a large-scale shotgun metatranscriptomics platform (total-RNA-seq) for host, bacterial, and viral profiling. We applied both technologies across 857 SARS-CoV-2 clinical specimens and 86 NYC subway samples, providing a broad molecular portrait of the COVID-19 NYC outbreak. Our results define new features of SARS-CoV-2 evolution, nominate a novel, NYC-enriched viral subclade, reveal specific host responses in interferon, ACE, hematological, and olfaction pathways, and examine risks associated with use of ACE inhibitors and angiotensin receptor blockers. Together, these findings have immediate applications to SARS-CoV-2 diagnostics, public health, and new therapeutic targets.

## Introduction

In March 2020, the World Health Organization (WHO) declared a pandemic of the coronavirus disease 2019 (COVID-19), an infection caused by severe acute respiratory syndrome coronavirus 2 (SARS-CoV-2) (He *et al.*, 2020). Since its start, more than three million of cases and more than two hundred of thousand deaths have been reported (https://coronavirus.jhu.edu), with an especially high burden in New York City (NYC). Since the presenting symptoms of COVID-19 can resemble those of common viral respiratory infections, a molecular diagnosis is required to reliably distinguish a SARS-CoV-2 infection from influenza and other respiratory illnesses (Guan *et al.*, 2020, Zhou *et al.*, 2020).

The current gold standard (qRT-PCR-based) approaches to SARS-CoV-2 molecular testing are largely limited to hospital or academic laboratories and reserved for the most severe cases, with limited accessibility to the general public. As a result, the true prevalence of SARS-CoV-2 in the population is mostly unknown, particularly among pre-symptomatic, mild symptomatic, or asymptomatic cases. Though several novel, scalable biotechnological approaches for viral testing have recently emerged (e.g., CRISPR-Cas12a (Broughton *et al.*, 2020) or CRISPR-Cas13 (Metsky *et al.*, 2020) on paper-based detection systems, or loop-mediated isothermal amplification (LAMP) (Tanner *et al.*, 2015, Zhang *et al.*, 2020, Yu *et al.*, 2020, Schmid-Burgk *et. al*., 2020), these have not been validated against gold-standard clinical assays or next-generation sequencing (NGS).

The lack of widely available, rapid SARS-CoV-2 diagnostics has fundamentally limited the public health approach to COVID-19, including hampering ability to implement contact tracing and accurate estimation of infection fatality rates. In addition, the persistence of SARS-CoV-2 across a range of surfaces (van Doremalen *et al.*, 2020) and hospital areas (Ong *et al.*, 2020) raises the specter of COVID-19 spread via fomite transmission. A key question is whether the environmental surface distribution of SARS-CoV-2 in high-touch, high-traffic areas (e.g. subways) may have driven its rapid emergence in certain regions (e.g., NYC). As yet, little information exists about RNA viruses in public areas.

Nonetheless, genomic viral surveillance resources, such as Global Initiative on Sharing All Influenza Data (GISAID), the Johns Hopkins University (JHU) Dashboard, and Nextstrain (https://nextstrain.org/ncov) (Gardy *et al.*, 2015, Dong *et al.*, 2020, Meyers *et al.*, 2020, Hadfield *et al.*, 2018) have enabled dynamic tracking of patient-derived viral evolution during the COVID-19 pandemic. These efforts still only cover a fraction (<0.5%) of documented cases, motivating ongoing viral profiling efforts. Unlike the results of simple binary diagnostic tests, full-length viral genome sequences can reveal patterns of strain evolution, temporal and geographic trajectories of viral spread, and correlations between specific genotypes and clinical features (e.g. disease severity, comorbidities, and viral load). These genomic epidemiology efforts have already played a crucial public health role in confirming community spread of SARS-CoV-2 in the USA (Zhao *et al.*, 2020, Gonzalez-Reiche *et al.*, 2020, Fauver *et al.*, 2020) as well as in China (Lu *et al.*, 2020). However, standard approaches to viral profiling (i.e. targeted sequencing) fail to provide information on either the host immune response or microbial co-infections, both which might impact the clinical presentation of COVID-19 and provide directions for therapeutic intervention and management.

To address these gaps in knowledge of disease and technology, we designed and validated a rapid reverse transcriptase LAMP (RT-LAMP) assay to detect SARS-CoV-2 infection from nasopharyngeal swab specimens and oropharyngeal swab lysates. We then integrated this assay into a large-scale host and viral profiling platform that combines targeted diagnostics with shotgun metatranscriptomics (**Figure 1a**). We applied this total RNA-seq platform to 857 samples from 735 confirmed or suspected COVID-19 cases at New York-Presbyterian Hospital-Weill Cornell Medical Center (NYPH-WCMC) and 86 environmental samples collected from high-transit areas in the NYC subway in early March 2020. Our results define genetic features of the NYC outbreak and implicate specific host responses in SARS-CoV-2 infection, including perturbations of the angiotensin converting enzyme (ACE) pathway.

**Figure 1.**
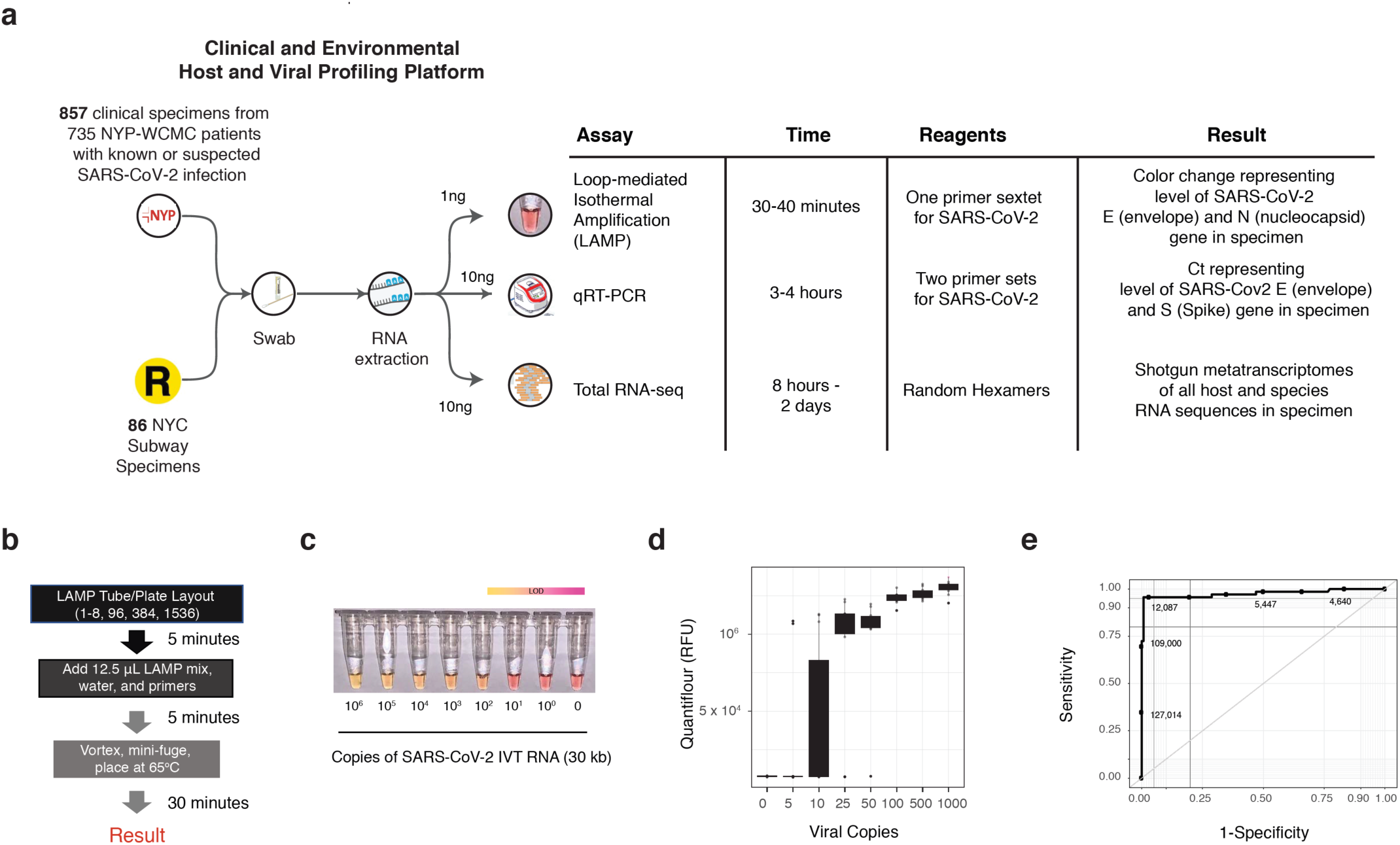
Sample Processing, the Loop-Mediated Isothermal (LAMP) Reaction and Synthetic RNA Validation. (a) Clinical and environmental samples collected with nasopharyngeal (NP) and isohelix swabs respectively, were tested with RNA-sequencing, qRT-PCR, and LAMP. (b) The test samples were prepared using an optimized LAMP protocol from NEB, with a reaction time of 30 minutes. (c) Reaction progress was measured for the Twist SARS-CoV-2 synthetic RNA (*MT007544*.*1*) from 1 million molecules of virus (10^6^), then titrated down by log10 dilutions. The colorimetric findings of the LAMP assay are based on a yellow to pink gradient with higher copies of SARS-CoV-2 RNA corresponding to a yellow color. The limit of detection (LoD) range is shown with a gradient after 30 minutes between 10 and 100 viral copies (lower right). (d) Replicates of the titrated viral copies using LAMP, as measured by QuantiFluor fluorescence. (e) The sensitivity and specificity of the LAMP assay from 201 patients (132 negative and 69 positive for SARS-CoV-2, as measured by qRT-PCR). Thresholds are DNA quantified by the QuantiFluor.

## Results

### Rapid, single tube detection of SARS-CoV-2

We developed a colorimetric assay to quickly detect and quantify SARS-CoV-2 viral load in patient and environmental samples, utilizing a set of six LAMP primers and simple single tube protocol (**Figure 1a-b**). Primers were designed to create two nested loops and amplify within the SARS-CoV-2 nucleocapsid gene (N gene), which enabled a 30-minute reaction workflow.

Related pathogens, high prevalence disease agents, and normal or pathogenic microbiota that are reasonably likely to be encountered in the clinical samples were evaluated to identify the percent homology between the primer sequences and these other organisms, and the probes were also designed to avoid known polymorphisms (see Methods).

To validate the assay, we first evaluated two synthetic RNAs (see Methods) whose sequences matched the viral sequences of patients from Wuhan, China and Melbourne, Australia (**Supp. Fig. 1**). The first control (MT007544.1) was used to test the analytical sensitivity via the limit of detection (LoD), titrated from 1 million molecules of virus (10^6^) down to a single copy, using serial log_10_ dilutions. The reaction output was measured at 0-, 20-, and 30-minute intervals (**Figure 1c, Supp. Figure 2a**) before the samples were heated to 95°C for inactivation. LAMP fluorescence correlated closely with SARS-CoV-2 RNA viral copies (**Figure 1d**), with an overlap of the median signal from negative controls at lower levels (0-10 total copies per reaction) of viral RNA (n=10). The LoD was found to be between 5–25 viral total copies for the dual primer, single-tube reaction (N gene and E gene), and this was then replicated to show a similar LoD on the second synthetic control from a patient from Wuhan (**Supp. Figure 2b, 3a**).

**Figure 2.**
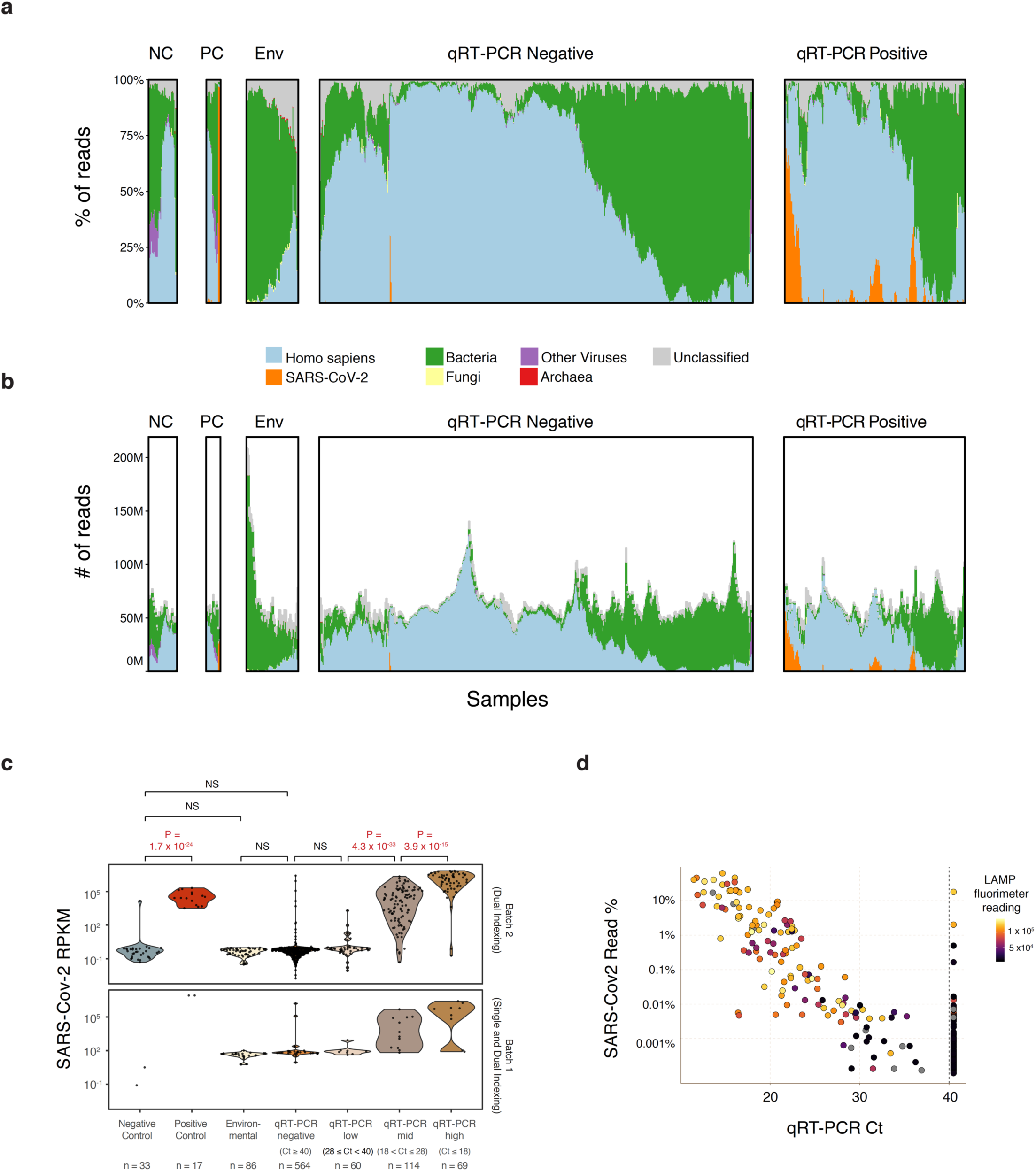
Full transcriptome profiles of SARS-CoV-2 Patients with NGS, qRT-PCR, and LAMP. (a,b) Read mapping to archaea (red), bacteria (green), fungi (yellow), human (blue), and SARS-CoV-2 (orange), and other viruses (grey), across the clinical controls (CN, CP), environmental samples, qRT-PCR negative, and qRT-PCR positive samples. (c) Clinical samples tested by qRT-PCR (Positive, n=255, or Negative, n=564) were sequenced and run through the LAMP assay. These results were compared to the buffer blanks (Negative Control, CN, n=33), Vero E6 cell extracts with SARS-CoV-2 infection (Positive Controls, CP, n=17), and Subway Samples (Environmental, Env, n=86). Read proportions are shown on the y-axis. (d) SARS-CoV-2 abundance, as measured with NGS and percentage of reads (y-axis) is compared to the Ct Threshold for qRT-PCR (x-axis), with lower Ct values representing higher viral abundance, and the LAMP reaction output (Fluorimeter values, black to yellow scale).

To optimize the LAMP assay for clinical samples, we first tested the reaction on a range of sample types, dilutions, and reaction volumes. We used a set of 201 samples from known or suspected COVID-19 cases that were tested for SARS-CoV-2 by a commercial qRT-PCR assay that has been the primary clinical test at NYPH-WCMC since early March. These comprised 69 nasopharyngeal (NP) swab samples that tested positive (qRT-PCR positives, Ct<40) and 132 samples that tested negative (qRT-PCR negative, Ct ≥ 40) (see Methods). qRT-PCR positive samples showed a much higher LAMP fluorescence than qRT-PCR negative samples, with superior performance observed at higher (10μL vs 5μL) input volumes (**Supp. Figure 3b**). qRT-PCR positive samples that failed to generate LAMP fluorescence were associated with lower viral genome abundance (high cycle threshold [Ct] value qRT-PCR). We obtained similar performance on bulk oropharyngeal swab lysate (**Supp. Figure 3c-e**), including increasing reaction sensitivity as a function of viral copy number. This lysate protocol required only a 30-minute lysis time for the oral collections and 30-minute LAMP reaction time. These results indicate robust performance of the LAMP assay across a broad range of purified nucleic acids as well as crude cellular lysates (see Methods).

**Figure 3.**
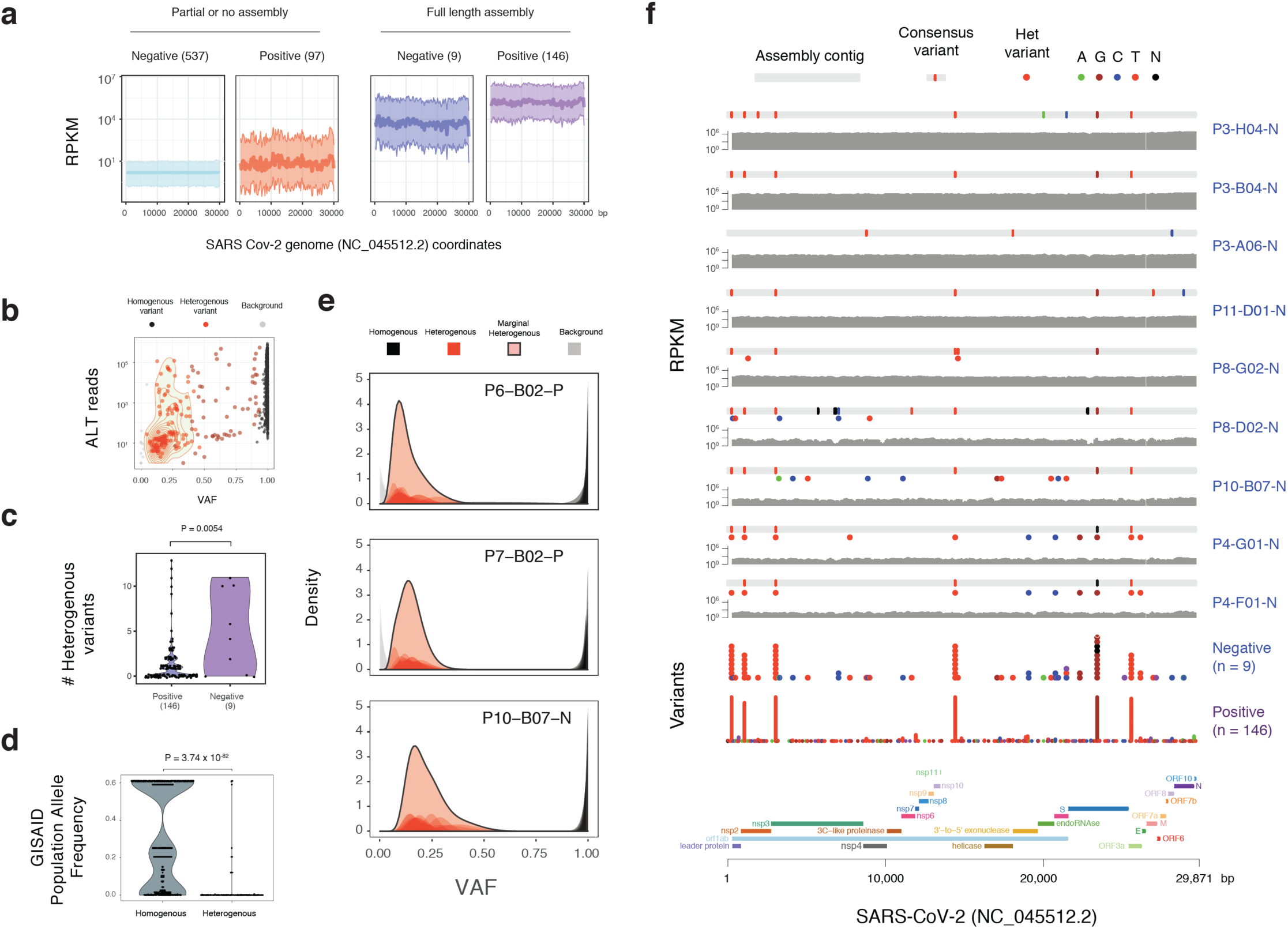
Viral genome assemblies and variants. (a) Clinical samples tested by qRT-PCR (Negative, N) were compared to RNA-seq coverage (Read per Kilobase per Million Reads, RPKM, y-axis) for the set of fully-assembled positive samples (n=146, 9) vs. those with partial or no assemblies (left, n=537, 97). (b) Variant allele frequencies (VAF, x-axis) for alternative alleles (y-axis) were calculated for all variants across viral strains, with heterogenous (het, 5%<x<95%, red) variants shown as well as homogenous (black) variants with VAF>0.95. (c) Proportion of het variants (y-axis) is shown for those tested by qRT-PCR and shown as negative or positive (x-axis). (d) The frequency of variants present in the GISAID global database of virus sequences (y-axis) is shown for variant types (x-axis). (e) The distribution and density of the VAF for three exemplar samples are shown relative to their variant type (top) (f) The assembled contigs (grey) the consensus variants (red dots) relative to the reference, and coverage (RPKM) are shown for a set of clinical samples, relative to the reference SARS-CoV-2 genome annotation (bottom), gene segments (colored bars), and histogram of variants from qRT-PCR positive and negative samples (variants y-axis).

Analysis of Receiver Operator Characteristic (ROC) curves revealed an optimal threshold of 11,140 QuantiFluor relative fluorescence units (RFUs), which yielded an overall sensitivity of 95.6% and specificity of 99.2% (**Figure 1e**). We observed higher LAMP sensitivity at higher viral loads, as determined by qRT-PCR Ct values (**Supp. Fig. 4**). The highest viral load (Ct <20) showed 100.0% sensitivity and 97.4% specificity, compared with the sensitivity at the lowest viral load at 80.0% (Ct >28, RFU below 7010). These same LAMP assay thresholds yielded consistent test positivity for synthetic RNA positive controls (Twist Biosciences) as well as clinical spike-in carrier RNAs (20/20) (**Supp. Fig. 4**), and consistent signal from clinical viral positives in Vero E6 cells (100.0%, 12/12) and blank clinical buffer negatives (100.0% 8/8).

**Figure 4.**
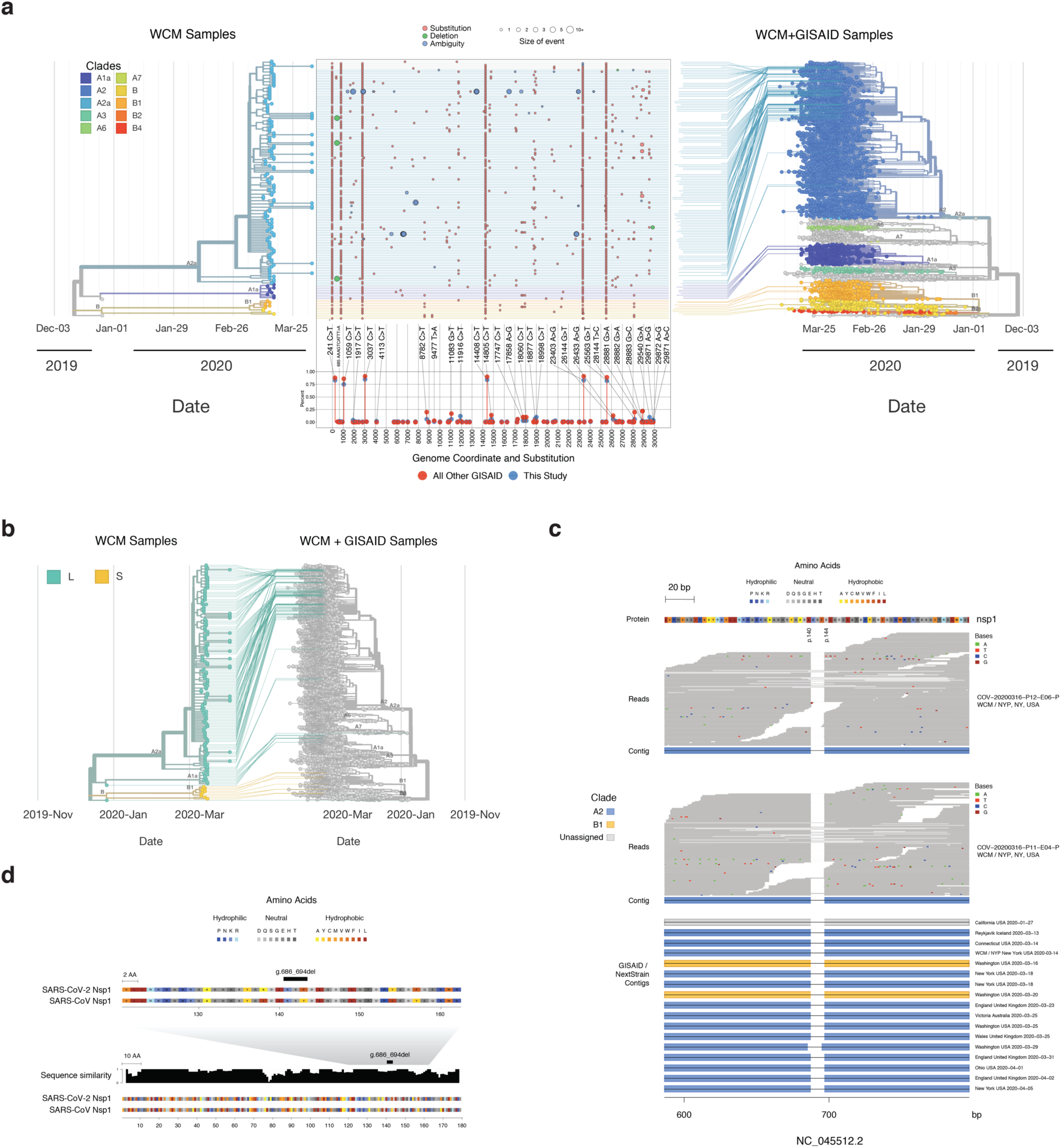
The mutational landscape SARS-CoV-2. (a) The phylogenetic placement of these SARS-CoV-2 samples is shown on the tree (left) and the global map of known SARS-CoV-2 genomes (right). Genetic variants called from the RNA-seq data (middle) show a range of variants that are distinct from the Wuhan reference strain, and the samples from this study, highlighted in blue, show enrichment for European and Asian alleles. The annotation track on the bottom shows variants called in these samples alongside all other GISAID samples. Samples that diverge from GISAID by more than 5% in either direction are annotated, including their coordinate and substitution event. (b) Proportion of the L (green) and S strain (yellow) are shown for the NYC viruses. Phylogeny of samples from this study on the left and total GISAID samples on the right, with a map of variants in this study’s samples in the middle, colored by event type and sized by number of nucleotides impacted. Annotation track on the bottom shows frequency of alternate alleles in this study and in the GISAID database. (c) The 9-bp deletion in ORF1b (NSP1 protein) that was detected in samples from three NYPH-WCMC patients was confirmed in the GISAID database (bottom tracks). Read-level support is shown in the top tracks for two of the variants, with aligned contigs visualized below. Assembly alignments for 17 additional cases harboring this deletion, including 1 additional case from this study are shown in the bottom track. (d) Alignment of SARS-CoV-2 to SARS-CoV NSP1 protein sequence is shown, with an enlarged view showing a sequence similarity track (normalized protein alignment score in 5 amino acid sliding window) and the 9-bp deletion region delineated.

### Shotgun metatranscriptomics platform for viral, microbiome, and host genomics

To provide orthogonal validation of our LAMP assay and further investigate the biological characteristics of qRT-PCR positive and negative specimens, we developed a shotgun metatranscriptomics platform utilizing total RNA-seq (RNA-sequencing with ribosomal RNA depletion) to profile all RNAs from patient and environmental swab specimens (**Figure 1a**). We sequenced 857 RNA-seq libraries (**Figure 2**) across 735 COVID-19 cases and controls (including the 201 tested with LAMP) to an average of 63.2M read pairs per sample. This spanned 243 qRT-PCR positive and 546 qRT-PCR negative samples, 17 positive controls (Vero E6 cells), and 33 negative controls (buffer). Replicate RNA-seq libraries from clinical samples were merged into a final set of 216 qRT-PCR positive samples and 519 qRT-PCR-negatives.

We aligned total RNA-seq reads to the human genome and NCBI reference databases (see methods), Kraken2 classification of non-human sequences revealed abundant bacterial or SARS-CoV-2 RNA alignments, with a lower proportion of sequences mapping to fungi, archaea, or other viruses (**Figure 2a**). Positive controls and clinical samples demonstrating qRT-PCR Ct values consistent with medium and high viral loads were significantly (p < 1 × 10^−16^) enriched in SARS-CoV-2 genome alignments (median reads per kilobase per million mapped reads, RPKM) relative to negative controls and qRT-PCR negative clinical samples (**Figure 2b**). A comparison of LAMP fluorescence, qRT-PCR Ct values, and total-RNA Seq RPKM across 201 clinical specimens demonstrated consistent estimates of SARS-CoV-2 viral abundance across these three technologies (R_seq_vs_Ct_ = −0.84, R_seq_vs_lamp_ = 0.82, R_lamp_vs_Ct_ = −0.80) (**Figure 2c, Supp. Figure 5a**), as well as RNA-seq extraction replicates (**Supp. Figure 5b**). These results provide additional validation for the LAMP SARS-CoV-2 assay on clinical samples.

**Figure 5.**
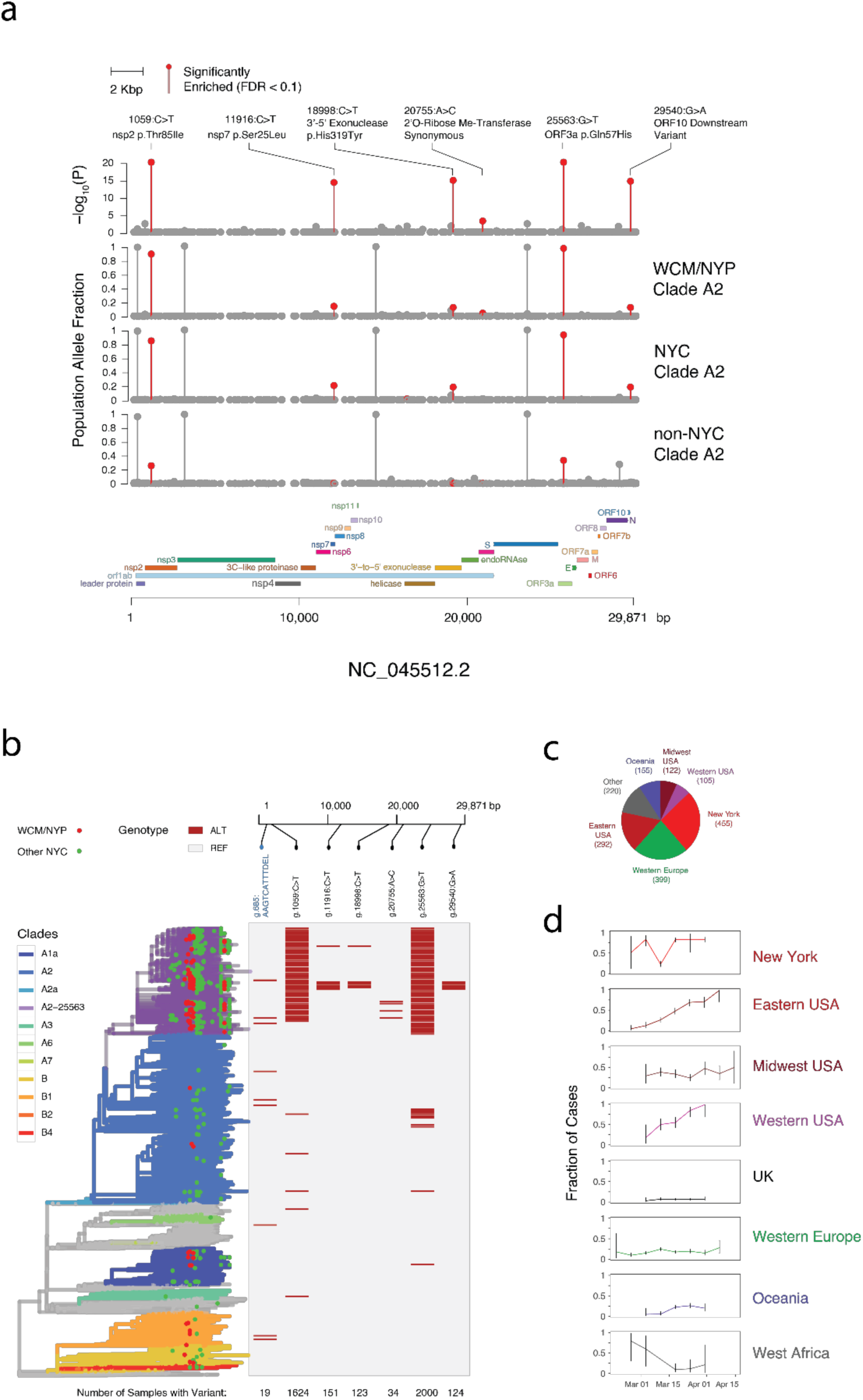
Variant and Subclade Analysis. (a) Six variant alleles were found to be significantly enriched by population allelic fraction within this set of 93 WCM/NYP cases as compared with non-NYC strains of Nextstrain clade A2 (1059:C>T P = 2.13 × 10^−47^, 11916:C>T P = 6.26 × 10^−15^, 18998:C>T P= 1.53 × 10^−15^, 20755:A>C P = 6.6 x10^−4^, 25563:G>T P = 7.92 × 10^−50^, 29540:G>A P = 2.68 × 10^−15^). These variants demonstrated similar PAF enrichment within all other NYC strains as compared to non-NYC clade A2 (1059:C>T P = 3.36 × 10^−146^, 11916:C>T P = 3.53 × 10^−74^, 18998:C>T P= 3.94x 10^−73^, 20755:A>C P = 7.71 × 10^−3^, 25563:G>T P = 1.45 × 10^−154^, 29540:G>A P = 2.07 × 10^−72^). (b) (left) Phylogenetic tree produced by the Nextstrain analysis with clade affiliations and nodes corresponding to WCM/NYP in red and other NYC cases in green. (right) occurrence of the six NYC-enriched alleles and the 9 nucleotide deletion across genomes. (c) Raw counts of cases present within this A2-25563 subclade demonstrated a predominance of European and North American cases, with Western Europe and New York together comprising the majority of strains. (d) Fraction of A2-25563 cases from each region of the world.

Analysis of total RNA-seq sequences also enabled mapping of likely co-infections and colonization with commensal species across both clinical positives and negatives in our sample set. We identified additional RNA viruses and organisms that distinguished high, medium, and lower viral load patients (**Supp Table 1, Supp. Figure 6**), with overall similarity observed across patients in the bacterial RNA fractions (**Supp. Figure 7a)**. However, other known respiratory viruses were observed in some of the COVID-19-positive patients (3.2%), including human coronaviruses 229E, coronaviruses NL63, influenza A, and human mastadenovirus and metapneumovirus (**Supp. Figure 7a)**. Most patients (209/216, 96.8%) that presented with the SARS-CoV-2 virus showed no evidence of other respiratory viruses, although bacterial mapping showed 102 patients with reads mapping to *Mycoplasma pneumonia* (relative abundance > 0.01%) (**Supp. Table 1**).

**Figure 6:**
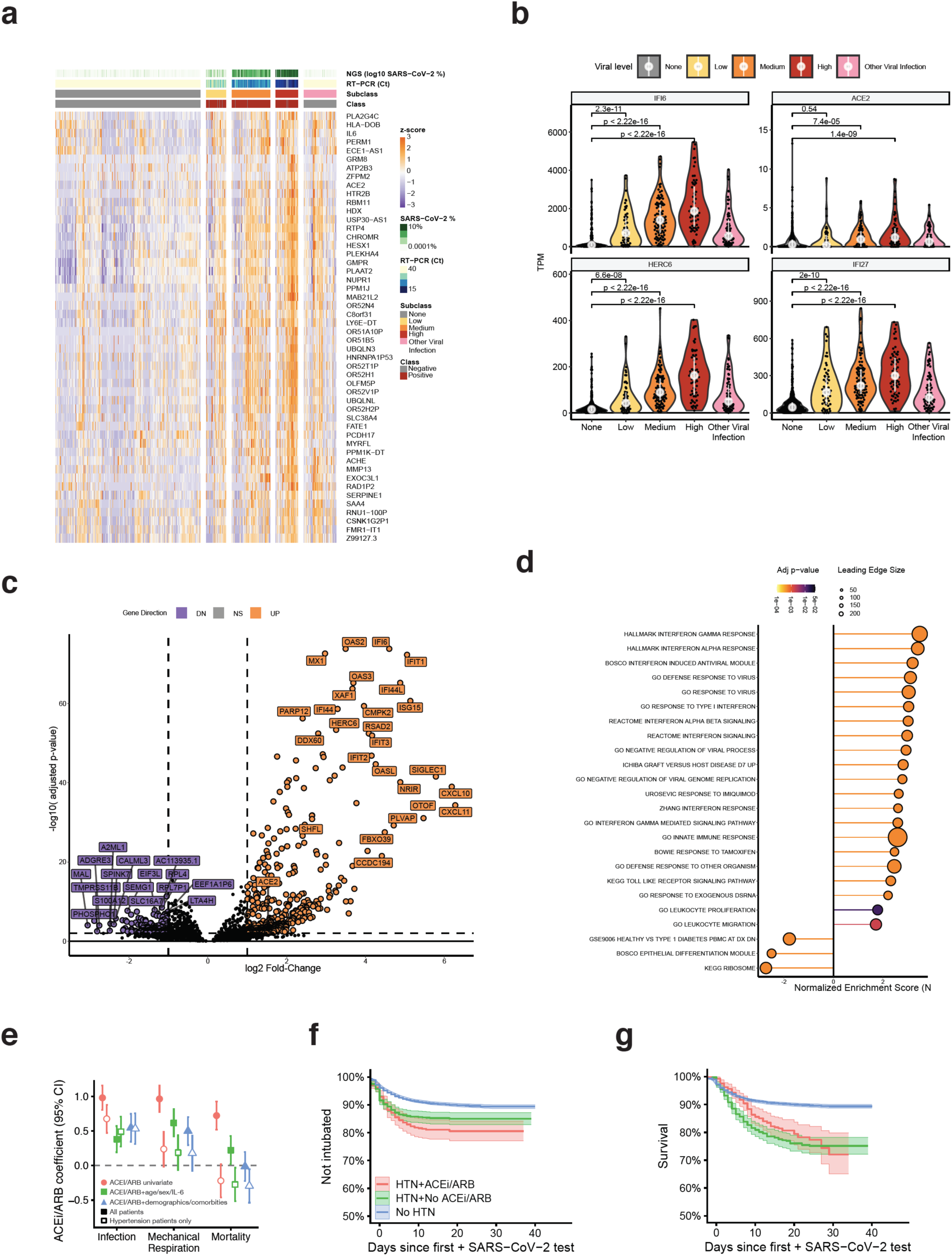
Host transcriptome responses and risk to SARS-CoV-2. (top row) Samples were quantified by a range of viral load, including RNA-seq (log10 SARS-CoV-2 % of reads), and qRT-PCR (Ct values) to create a three-tier range of viral load for the positive samples (right) compared to the clinically-annotated negative samples (class, red or grey) and those samples with other viral infections that were SARS-CoV-2 negative by qRT-PCR. (b) The differentially expressed genes of qRT-PCR positive patients compared to qRT-PCR patients showed up-regulated (orange) genes as well as down-regulated (purple) genes. (b) Up-regulated genes, with boxplots across all samples, include *IFI6, ACE2, SHFL, HERC6, IFI27*, and *IFIT1*, based on data from (c), which is the total set of DEGs. The full set is shown in an intersecting heat map, with a core set of up-regulated genes (orange) distinct form the set of down-regulated genes (purple), compared to genes that are not significantly differently expressed (grey) in any comparison (DESeq2, q-value <0.01, |logFC| >0.58). (d) GSEA enrichment of significant pathways, with color indicating statistical significance and circle size the number of genes on the leading edge (e) Regression coefficients for variables indicating exposure/history of exposure to ACEI/ARBs inhibitors for each of the three cohort comparisons: (left) test outcome in a cohort of patients suspected of SARS-CoV-2 infection, (middle) requirement of mechanical respiration in patients who tested positive, (right) mortality in patients who tested positive. Univariate analyses are shown as red circles. The green triangles coefficients are when correcting for age, sex, and baseline IL-6 levels take upon admission. The blue squares are from a model that includes age, sex, and IL-6 as well as comorbidities including CAD/CHD, diabetes, being overweight, or obesity, and self-reported race and ethnicity. Additionally, open markers indicate the same analyses but restricted to only patients with clinically diagnosed hypertension. (f) Curves for patients requiring mechanical respiration (identified by intubation procedure notes) as a function of drug class (g) and survival, conditioned on hypertension status and ACEI/ARB exposure status.

### Environmental sampling of SARS-CoV-2 in the NYC subway

Having validated a rapid LAMP assay and shotgun metatranscriptomics platform, we next investigated the environmental distribution of SARS-CoV-2 in high-transit areas of the NYC subway at the beginning of the NYC pandemic. We collected 62 samples from handrails, kiosks, and floors in Grand Central and Times Square subway stations between March 6 and 13th, 2020 and generated 86 RNA-seq libraries. Each sample was collected using a sterile and DNA/RNA free swab, following the MetaSUB sampling protocols for nucleic acid stabilization (Danko *et al.*, 2020).

To further investigate these 86 environmental samples for the presence of SARS-CoV-2, we generated total RNA-seq libraries and investigated the distribution of non-human sequences. These samples demonstrated a mix of fungal and archaeal species that was consistent with underground subway origin (Danko *et al.*, 2020) (**Supp. Table 1**). However, we did not observe significant counts or proportions (**Figure 3a**) of SARS-CoV-2 reads, which was particularly clear with dual-index library preparations (**Supp. Figure 8**). However, a broad range of other bacterial and viral species was found, including a large set of phages (e.g. *Streptomyces* phage VWB), desiccation-tolerant bacteria (e.g *Deinococcus radiodurans*), and more abundant bacterial and archaeal RNA than the clinical samples (**Supp. Table 1**). Of note, these were checked against a database of putative false positives (**Supp. Table 2**), which was created from *in silico* fragmentation of the SARS-CoV-2 genome and mapping against the same database. Taken together, these results indicate that high transit surfaces were not likely to harbor significant levels of SARS-CoV-2 in the early phases of the NYC epidemic.

### SARS-CoV-2 assemblies from shotgun metatranscriptomes

The abundance of SARS-CoV-2 alignments from total RNA-seq was sufficient to provide >10x coverage of the viral genomes and yield high quality, full-length assemblies for 155 samples, including 146 (67.6%) of qRT-PCR positive patient samples (**Figure 3a, Supp. Figure 6**). The 97 qRT-PCR positive samples that yielded partial or no assembly demonstrated low or uneven genome-wide read depth, though these were increased on average relative to qRT-PCR negatives. We also generated full length SARS-CoV-2 assemblies for 9 samples that were found negative by qRT-PCR (**Figure 3a**). Each of these demonstrated a high RPKM viral load, an abundance of reads evenly covering the SARS-CoV-2 genome, and high (>0.5) variant allele fractions (VAFs) of SNV’s commonly seen in positive samples, as well as a recent build of GISAID (**Figure 3b**). Apart from these common variant sites, we did not find additional recurrent mutations that might explain why these cases were not detected by qRT-PCR (e.g. primer site mutation).

When examining the viral genomes, we identified a total of 1,147 instances of 165 unique variants across 155 assemblies, including 1,143 single nucleotide variants (SNVs) and 4 deletions. Variant calling of assemblies and RNA-seq reference alignments showed most (87.5%) variants with VAFs >0.95 and high (>100) numbers of variant-supporting reads. Analysis of VAF posterior probability distributions identified a subset of alleles whose VAFs were confidently (probability > 0.95) above 0.05 but below 0.95. We labeled these alleles as “het” (heterogenous) variants and refer to the remaining (high VAF) consensus assembly variants as “homogeneous.” Many of these het variants were associated with robust read support despite frequently having VAFs below 0.5 and not being incorporated into the consensus assembly sequence. A subset of samples (11/155) samples harbored at least 5 het variants, with a significant enrichment (P = 0.0054) in qRT-PCR negative cases (**Figure 3c**). Analyzing the population allele frequencies (PAF) of these het variants across GISAID revealed the vast majority (91%) to be rare (PAF<0.1) or even private to that sample, indicating that these were unlikely to be due to cross contamination (**Figure 3d**).

We then asked whether het variants might indicate the presence of a viral subclone, arising either through co-infection or intra-host adaptation. Analyzing het-rich (>5 het variants) samples, revealed that set of het variants observed in a single sample showed similar VAFs, yielding clear peaks when summing their individual posterior probability densities (**Figure 3e)**. These results were consistent with the presence of a viral subclone at some fraction *f* corresponding to the modes of each of these VAF distributions (0.1-0.2). Interestingly, each of the three samples shown in **Figure 3e** also harbored a clear peak of homogenous variants with VAFs near 1. These results suggest that the viral subclone is not the result of a coinfection by two viruses, which would yield a second peak at 1-*f* at every major clone variant that has the reference allele in the minor subclone. Het variants were found to occur at common sites of consensus variants across samples, however, rare private hets were also present (**Figure 3f**).

### Global phylogenetic relationship of NYC strains

We next integrated 141 full-length NYPH-WCMC sequences with 7,806 SARS-CoV-2 sequences obtained from a recent GISAID build (downloaded on 4/20/2020) into a maximum likelihood phylogenetic tree using the Nextstrain algorithm (Hadfield *et al.*, 2018) (**Figure 4a**, see Methods). This integration resulted in 141/155 NYPH-WCMC sequences integrated into the tree. Analyzing worldwide groupings annotated by the Nextstrain database, we found a high proportion (>87%, 123/141) of incorporated samples were associated with A2, a large Western European-derived clade that comprises nearly half of the sequences in GISAID, which agrees with recently published findings (Gonzalez-Reiche *et al.*, 2020). In addition, we found a minority of samples from clades with mixed Asian and European origins (A1a, 6% of cases; B1, 4% of cases; B, 3% of cases). Among these, we identified a clear predominance (94.1%, 146/155) of the “L” strain, which is defined by a reference base at position 28143 (ORF8) and previously associated with severe cases in Wuhan (Tang *et al.*, 2020) (**Figure 4b**).

Among the A2 cases in our cohort, we found a recurrent 9 bp in-frame deletion (g.686_694del) in the gene encoding non-structural protein 1 (NSP1). NSP1 is a putative SARS-CoV-2 virulence factor that is highly divergent across betacoronaviruses (Narayanan *et al.*, 2015). This deletion was assembled in 3 of 93 NYPH-WCMC samples, in which it showed robust read support with near >0.95 VAF across thousands of high-quality alignments to the locus (**Figure 4c**). An identical deletion was observed in 15 other samples in the GISAID database, spanning samples from England, Iceland, and Canada, as well as three additional cases from New York. Among these, 12 of 15 were labeled as part of clade A2 by Nextstrain, while two (Washington state) cases were associated with a distant clade B1 (**Figure 4d**). This deletion removes three amino acids (p.141_143KSD), including a residue that is variant (143Y>F) relative to the SARS-CoV-1 genome (**Figure 4c**). Interestingly, a 16th case (Washington, clade A2, **Figure 4c**) showed a related but not identical 9 bp in-frame deletion (g.686_694del) that removes the same three residues. These residues occur in a conserved portion of the C-terminal region of NSP1, which has been linked to host chemokine dysregulation and translational inhibition in SARS-CoV (Narayanan *et al.*, 2015).

### A novel SARS-CoV-2 subclade enriched in NYC cases

Given the predominance of clade A2 among NYC cases, we next asked whether specific variants were enriched in NYC A2 samples relative to other regions’ A2 samples. Comparing variant frequencies between A2 cases in our data and non-NYC cases in GISAID, we identified six enriched loci with FDR<0.1 (Fisher’s exact test, one-tailed) that were over-represented in NYC A2 samples (**Figure 5a**). All six of these variants were significantly enriched (Bonferroni-adjusted p-value <0.05) in a second analysis comparing all other cases from NYC in GISAID vs. the rest of the world (**Figure 5a**). Visualizing the most significant of these variants across the Nextstrain derived phylogenetic tree (**Figure 5b**) demonstrated that the highest frequency of these variants (25563G>T), defined a novel subclade of A2, which we call A2-25563. This subclade comprised 84.4% (572/678) of (NYPH-WCMC and GISAID) NYC samples, and 92.7% (572/617) of the A2 portion of NYC cases (Sinai and NYU). Interestingly, while all three of the NYPH-WCMC A2 cases harboring the aforementioned NSP1 9-bp deletion were in A2-25563, many of the remaining cases were distributed across other subclades. Unlike the SNVs used to define A2-25563, this deletion appeared to be highly polyphyletic with 7 unique phylogenetic clusters associated with the deletion across a variety of clades (**Figure 4c, Figure 5b**).

Investigating the geographical and temporal distribution of non-NYC samples in A2-25563, we found a global distribution with likely origin in Western Europe (**Figure 5c, 5d**). The first noted case of A2-25563 in GISAID was in Belgium on 2/21/2020, but since then, this putative subclade has represented a minority (20%) of Western European cases. While the prevalence of A2-25563 in NYC appeared to be consistently high across sampling dates, it constituted only a moderate proportion of cases from the Western and (non-New York portion of the) Eastern USA (<50%) in early March (**Figure 5d**). Strikingly, we found that of the prevalence of the A2-25563 clade across Eastern and Western United States steadily and significantly increased through late March and early April, while remaining stable in the Midwest. Among the early locations showing high fractions of A2-25563 was West Africa, which has steadily decreased in prevalence since early March (**Figure 5d**). The SNVs defining A2-25563 subclade were non-synonymous variants targeting genes encoding the non-structural protein 2 (NSP2), the viral replicase, and ORF3a, a poorly characterized SARS-CoV-2 protein with putative roles in inflammation (Siu *et al.*, 2019). This subclade defining ORF3a site (p.Gln57His) is distinct from that which defines the previously characterized L strain (p.Leu84Ser) (Tang *et al.*, 2020).

### Defining the SARS-CoV-2 host transcriptome

We leveraged the comprehensive nature of the total RNA-seq profiles to better understand the host transcriptome during SARS-CoV-2 infection. Cell proportion analyses using the MUSIC deconvolution algorithm showed enriched proportions of cell types spanning goblet, ciliated airway, and epithelial cells across all samples (**Supp. Figure 9**). Differentially expressed genes (DEGs) associated with SARS-CoV-2 infection were calculated using limma voom and DESeq2 (see Methods). Overall, 757 significant DEGs (q<0.01, >1.5-fold change) were found the qRT-PCR positive vs. negative samples (**Supp. Table 3)**, spanning 350 up-regulated DEGs and 407 down-regulated DEGs, and a total of 8851 unique genes (q<0.01, >1.5-fold change) across different subgroups including a subset that were distinct from other detected viral respiratory infections (**Figure 6a, Supp. Figure 12**). These spanned viral response pathways, innate immune response, and interferon signaling (**Figure 6b,d, Supp. Figure 10a**), but also included iron, olfaction, calcium, and ribosome dysregulation.

Differentially expressed host genes indicated a wide range of antiviral responses, including a common interferon response across all ranges of viral levels, which was significantly higher when compared to SARS-CoV-2 negative samples that harbored other respiratory viruses (**Figure 6a,b**). Notably, host cells showed an increase in angiotensin converting enzyme 2 (*ACE2)* expression (**Figure 6b**) (p-value=1.4× 10^−9^), which is the SARS-CoV-2 cellular receptor (Hoffmann *et al.*, 2020). This critical gene for viral entry (Sungnak *et al.*, 2020) exhibited a dose-dependent expression concomitant with the higher levels of SARS-CoV-2 virus, along with IFI27 (Interferon Alpha Inducible Protein 27, p < 2.2× 10^−16^) and IFI6 (Interferon Alpha Inducible Protein 6, p < 2.2× 10^−16^). The DEGs also included HERC6 (HECT Containing E3 Ubiquitin Protein Ligase Family Member 6), which aids Class I MHC-mediated antigen processing and Interferon-Stimulated Genes (ISGs) (**Figure 6b,c,d**), underscoring the impact of the virus on these cells’ immune response (Oudshoorn *et al.*, 2012). Also, a subset of cytokines (CXCL10, CXCL11, CCL8) showed the highest spike of expression in the higher viral load sub-group (**Supp. Figure 10**), matching previous results from animal models and infected cells (Blanco-Melo *et al.*, 2020). The host transcriptome that exhibited the greatest amount of DEGs were those with the highest viral titer (**Figure 6a**).

Down-regulated genes and those with a negative enrichment score (NES) were functionally distinct (**Figure 6d**). This included a significant decrease in gene expression for the olfactory receptor pathway genes (q-value =0.0005, **Supp. Table 4**), which is consistent with a COVID-19 phenotype wherein patients lose their sense of smell. Other down-regulated genes included the transmembrane serine protease *TMPRSS-11B*, which regulates lung cell growth (Updegraff *et al.*, 2018) and *ALAS2*, a gene which makes erythroid ALA-synthase (Ajioka *et al*,. 2006) that is found in developing erythroblasts (red blood cells). ALA-synthase plays an important role in the production of heme TRIM2 E3 ubiquitin ligase induced during late erythropoiesis, which indicated a connection to hematological and iron (heme) regulation during infection (**Figure 6d**). Accordingly, genes in a related biological network were significantly enriched based on Gene Ontology (GO) pathways for iron regulation (q-value 0.04, **Supp. Table 4**). Both the up-regulated and down-regulated gene expression differences in were distinct from those of house-keeping genes (**Supp. Figure 11**), which stayed mostly stable during infection.

### ACE inhibitor/angiotensin receptor blocker usage correlates with COVID-19

Finally, given our observation of increased *ACE2* gene expression in patients with high SARS-CoV-2 viral load, we then investigated the interplay of receiving pharmacologic angiotensin converting enzyme inhibition (ACEI) or angiotensin receptor blockers (ARBs) for hypertension and clinical features of COVID-19 disease. Since *ACE2* expression can be increased in patients taking ACEIs and ARBs (Agata *et al.*, 2006), the observed correlation of viral titer with *ACE2* expression may be attributed to the pre-infection use of such inhibitors, which is common in older patients and those with comorbidities (Fang *et al.*, 2020). To address this, we analyzed an observational cohort of 8,856 patients with suspected SARS-CoV-2 infection from New York Presbyterian Hospital-Columbia University Irving Medical Center (NYPH-CUIMC) for their usage of ACEIs and ARBs (4,829 who tested positive). We found that use of ACEIs/ARBs was strongly associated with testing positive in patients suspected of SARS-CoV-2 infection (Odds Ratio, OR=2.7, and 95% Confidence Interval, CI=2.2-3.94, p=7.43E-28, **Figure 6e**). This result was consistent when corrected for age, sex, and IL-6 (OR=1.5 CI:1.2-1.8), when corrected for comorbidities (**Table 1**) (OR=1.754 CI:1.4-2.0) and when only considering patients with hypertension (OR=1.8, CI: 1.4-2.3). For comparison, we repeated this analysis with calcium channel blockers (CCBs), another class of anti-hypertensives, and found no association between CCB exposure and infection. We found that patients with hypertension who take ACEIs and ARBs have higher infection rates than patients with hypertension on CCBs (OR=1.5 CI: 1.2-2.0, p=0.0011) (**Table 2**).

**Table 1.** Baseline characteristics of SARS-CoV-2 suspected cohort. General population refers to a comparison cohort of individuals administered drugs at NYPH-CUIMC between January 1, 2019 – September 24, 2019 who were not later tested for SARS-COV-2 infection.

| Variable | General | Tested | COV+ | COV+/Intubated | COV+/Died |
| --- | --- | --- | --- | --- | --- |
| N (% of tested) | 90989 | 8856 (100%) | 4829 (54.5%) | 534 (6%) | 528 (6%) |
| ACEI | 2201 (2.4%) | 408 (4.6%) | 310 (6.4%) | 78 (14.6%) | 57 (10.8%) |
| ARB | 2113 (2.3%) | 400 (4.5%) | 295 (6.1%) | 70 (13.1%) | 65 (12.3%) |
| ACEI and ARB | 163 (<0.1%) | 28 (0.3%) | 22 (0.5%) | 7 (1.3%) | 4 (0.8%) |
| Median age (50%) | 34.6 (27.5-42.2) | 57.2 (38.3-71.8) | 62.4 (47.8-75) | 65.4 (54.4-74.8) | 78.2 (69.9-87) |
| Male | 31228 (34.3%) | 4001 (45.2%) | 2546 (52.7%) | 336 (62.9%) | 309 (58.5%) |
| Black/African-American | 1739 (1.9%) | 2047 (23.1%) | 1229 (25.5%) | 125 (23.4%) | 95 (18%) |
| Caucasian | 108 (0.1%) | 2587 (29.2%) | 1249 (25.9%) | 127 (23.8%) | 141 (26.7%) |
| Asian or Pacific Islander | 1564 (1.7%) | 237 (2.7%) | 106 (2.2%) | 4 (0.7%) | 4 (0.8%) |
| Other race | 79 (0.1%) | 2134 (24.1%) | 1250 (25.9%) | 156 (29.2%) | 165 (31.2%) |
| Missing race | 140 (0.2%) | 1851 (20.9%) | 995 (20.6%) | 122 (22.8%) | 123 (23.3%) |
| Hispanic/Latino | 13875 (15.2%) | 2811 (31.7%) | 1704 (35.3%) | 260 (48.7%) | 251 (47.5%) |
| Non-Hispanic/Latino | 23537 (25.9%) | 3576 (40.4%) | 1811 (37.5%) | 151 (28.3%) | 156 (29.5%) |
| Other ethnicity |  | 95 (1.1%) | 61 (1.3%) | 7 (1.3%) | 13 (2.5%) |
| Missing ethnicity | 53577 (58.9%) | 2374 (26.8%) | 1253 (25.9%) | 116 (21.7%) | 108 (20.5%) |
| Hypertension | 15298 (16.8%) | 2200 (24.8%) | 1392 (28.8%) | 243 (45.5%) | 310 (58.7%) |
| CAD/CHD | 1978 (2.2%) | 831 (9.4%) | 511 (10.6%) | 92 (17.2%) | 137 (25.9%) |
| Diabetes | 7239 (8.0%) | 1340 (15.1%) | 867 (18%) | 163 (30.5%) | 203 (38.4%) |
| Overweight | 12878 (14.2%) | 923 (10.4%) | 560 (11.6%) | 95 (17.8%) | 101 (19.1%) |
| No RF | 64695 (71.1%) | 6223 (70.3%) | 3212 (66.5%) | 264 (49.4%) | 198 (37.5%) |
| Median IL-6 (50%) | - | 67.8 (29.3-152.9) | 70.2 (30.8-154.4) | 152.8 (72.9-278.1) | 146.1 (81.2-284.8) |

**Table 2.**
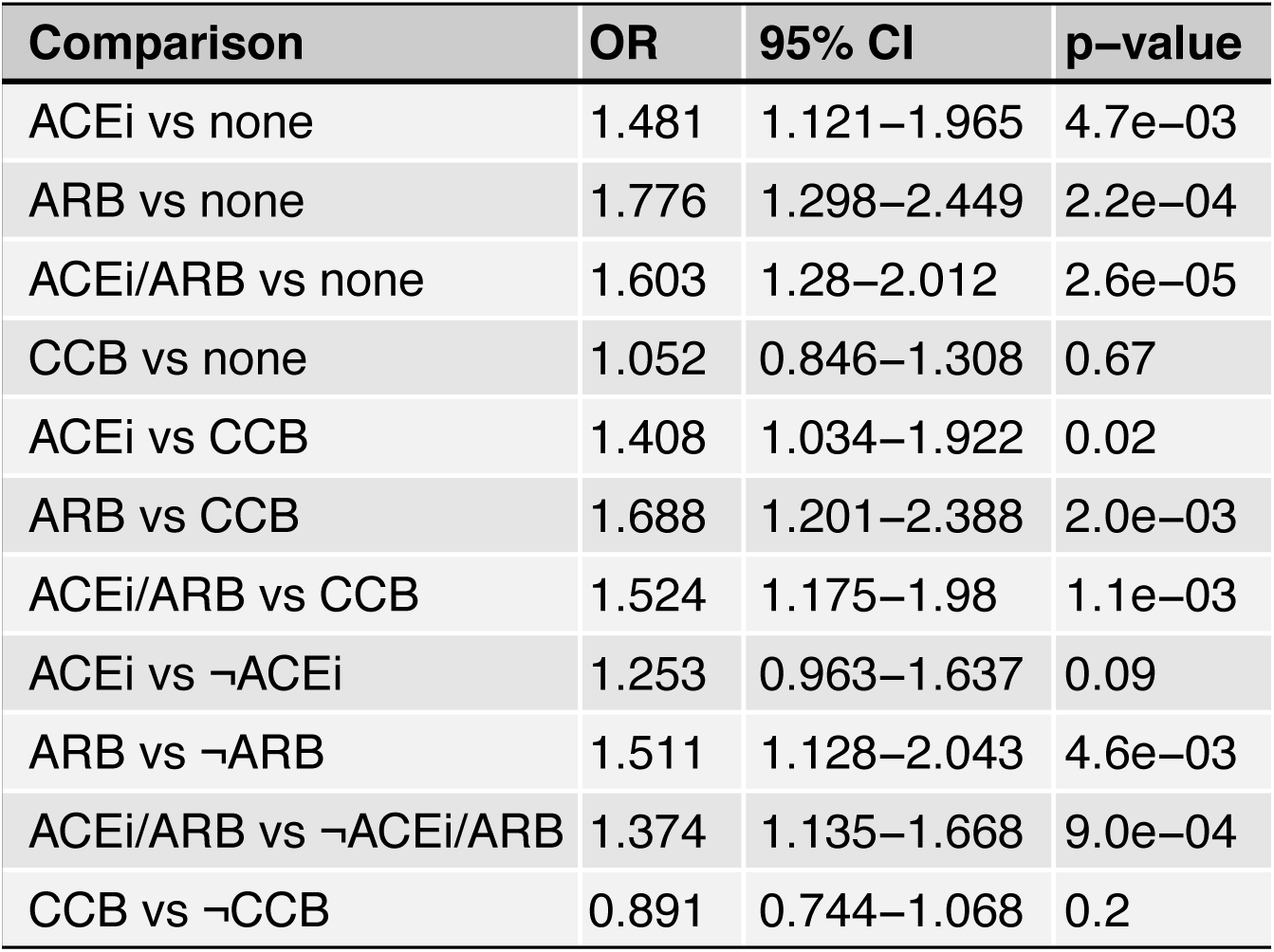
Comparison between ACE/ARB exposure and CCB exposure. To evaluate if evidence of hypertension treatment is confounding association between ACE/ARB exposure and infection. These analyses were completed using only patients with hypertension. In each test the ACE/ARB groups showed significantly greater rates of infection. CCB exposure was not found to be associated with infection.

| Comparison | OR | 95% CI | p-value |
| --- | --- | --- | --- |
| ACEi vs none | 1.481 | 1.121–1.965 | 4.7e–03 |
| ARB vs none | 1.776 | 1.298–2.449 | 2.2e–04 |
| ACEi/ARB vs none | 1.603 | 1.28–2.012 | 2.6e–05 |
| CCB vs none | 1.052 | 0.846–1.308 | 0.67 |
| ACEi vs CCB | 1.408 | 1.034–1.922 | 0.02 |
| ARB vs CCB | 1.688 | 1.201–2.388 | 2.0e–03 |
| ACEi/ARB vs CCB | 1.524 | 1.175–1.98 | 1.1e–03 |
| ACEi vs ¬ACEi | 1.253 | 0.963–1.637 | 0.09 |
| ARB vs ¬ARB | 1.511 | 1.128–2.043 | 4.6e–03 |
| ACEi/ARB vs ¬ACEi/ARB | 1.374 | 1.135–1.668 | 9.0e–04 |
| CCB vs ¬CCB | 0.891 | 0.744–1.068 | 0.2 |

We found a range of effects when investigating the relationship between ACEI/ARB usage and morbidity and mortality. In an uncorrected univariate analysis, ACEI/ARB usage conferred an increased risk of intubation and mortality for SARS-CoV-2 positive patients (intubation: HR=2.8 CI:2.3-3.5, 6.99E-24) (**Figure 6e** and **Figure 6f**) and mortality: HR=2.2 CI:1.8-2.7, p=3.18E-14 (**Figure 6e** and **Figure 6g**). These results hold when correct for age, sex, and IL6 (**Figure 6e**). However, when correcting for comorbidities, only intubation was significant (HR=1.7 CI:1.4-2.2, p=5.91E-07). Further, in the analysis where only patients with hypertension were included, we found modest evidence for both (i) a risk of intubation in the univariate model (HR=1.4 CI:1.1-1.8, p=0.02) and (ii) a protective effect of ACEI/ARBs for mortality in the covariate models (HR=0.8 CI:0.6-1.0, p=0.03).

Additionally, we confirmed previously reported risk factors for both mechanical respiration and mortality. For requirement of mechanical respiration, we found significant risk effects from male sex (p=2.76E-03), diabetes (p=2.64E-02), and IL-6 (p=1.07E-121). In addition, for mortality we found significant effects from age (p=1.10E-86), male sex (p=1.98E-04), diabetes (p=6.04E-04), being overweight (p=3.86E-03), and IL-6 (p=8.68E-23). Finally, in a post-hoc comparative analysis between specific ACEIs, we found no significant differences in infection rates, intubation, or mortality with any single ACEI drug over another (**Supp. Figure 13)**. In a similar analysis on ARBs, we found that olmesartan was associated with an increased risk of mortality (HR=2.3 CI: 1.1-4.9, p=0.03, N=28) in the hypertension-only analysis. We found no risk in the larger cohort nor for any other single ARB (**Supp. Figure 13**, with complete modeling results are provided in **Supplemental Tables 6-9**).

## Discussion

During a large-scale pandemic with exponential spread, such as COVID-19, scalable methods for diagnosis and screening are crucial for both mitigation and containment (Lan *et al.*, 2020, Liu *et al.*, 2020). While hospital-grade, core lab devices can achieve massive throughput (thousands of samples per day), a key limitation of these assays is accessibility of testing facilities to patients, the logistics of sample transport, and timely test reporting. These limitations become even more stark in the context of widespread quarantines and nationwide lockdowns, where requiring patients to travel (even for viral testing) incurs significant personal and public health risks.

The most urgent diagnostic need in this situation is for scalable rapid point-of-care tests that can be potentially implemented in the home. Our validation of a rapid one-tube, dual-primer colorimetric SARS-CoV-2 assay with both qRT-PCR and total RNA-seq provides a potential solution to this problem. Further work will be needed to assess whether this LAMP assay can detect the presence of SARS-CoV-2 at even lower (but clinically relevant) viral concentrations in specimen types that are less cumbersome to collect than naso/oropharyngeal swabs (e.g. saliva, stool). As we demonstrate, this LAMP SARS-CoV-2 assay can be also applied for environmental sampling, which may be crucial in the containment and recovery phases of this pandemic. Specifically, LAMP positivity may quickly indicate if an area is infectious and a negative result (with appropriate confirmation) will possibly represent a lower risk. Indeed, these tools and methods can help create a viral “weather report” if broadly used and partnered with continual validation.

Total RNA-sequencing data enabled a complete genetic map of the viruses in a significant subset of our samples. Remarkably, we found 9 qRT-PCR negative cases among the 155 cases that had sufficient reads to assemble the SARS-CoV-2 genome *de novo*. Though these likely qRT-PCR false negatives could not be attributed to specific sequence changes (e.g. primer site mutations), their high frequency underscores the limitation of “gold standard” qRT-PCR approaches for SARS-CoV-2 detection. These results further highlight the need for open-source primer design (the RT-PCR methods for this FDA-approved test are proprietary), so these assays can be updated as a more granular picture of strain diversity and evolution is obtained through worldwide sequence efforts.

Notably, our phylogenetic analysis nominates an A2 subclade (A2-25563, defined by 25563G>T) which comprises the majority of known NYC samples, including those sequenced outside of this study (Gonzalez-Reiche *et al.*, 2020). Though remaining NYC cases show a wide distribution across all identified clades, the predominance (>80% in NYC) of such a narrowly defined set of sequences within NYC from a rare (≤20%) Western European subclade is striking. The steady increase of A2-25563 prevalence in other regions in the Eastern and Western USA (but not Midwest) through late March and early April may be consistent with emigration patterns of NYC residents who left the city during the lockdown. The dominance of the proposed A2-25563 subclade in NYC suggests either (1) a very early introduction by a single patient harboring A2-25563; or (2) multiple A2-25563 founder events; or (3) disproportionate community transmission of strains within this subclade. The first possibility may be potentially linked to expansion among cases in the New Rochelle cluster, though additional genotyping and/or metadata aggregation from those sequenced cases would be required to assess this hypothesis. The third possibility could be consistent with A2-25563 harboring differential fitness with respect to transmissibility or virulence relative to other A2 viruses.

Future studies correlating viral genotypes with patient outcomes in larger cohorts will be necessary to determine whether any of these A2-25563 associated variants functionally influence viral transmission or disease severity. The polyphyletic pattern and convergent evolution pattern observed with NSP1 p.141_143KSD, including clades outside of A2, raises the possibility that this variant arose multiple times and may be under positive selection. Given the small number of these variants observed in our analysis (14), larger and more statistically powered datasets will be required to evaluate this hypothesis. The VAF distribution in patients harboring het variants suggests that intrahost diversification may lead to the development of SARS-CoV-2 subclones in a subset of COVID-19 cases. The relatively large numbers of these het variants (>10) presumably arising during the course of a single infection rising to a reasonably high oligoclonal VAFs (0.1-0.2) suggests that this intrahost diversification may be rapid and also associated with positive selection.

Our results demonstrate that distinct host transcriptional programs are activated during viral infection of the naso-/oropharynx with SARS-CoV-2. This includes upregulation of specific interferon pathway genes (*SHFL, IFI6, IFI27*, and *IFIT1*). that have been previously associated with the innate antiviral host immune response against other positive-strand RNA viruses (e.g. hepatitis C, Dengue virus). These results provide clinical relevance for recently published results from animal and cellular models of SARS-CoV-2 (Blanco-Melo *et al.*, 2020). Our analyses also implicate expression perturbations of the ACE pathway in SARS-CoV-2 host response, including *ACE2*. ACE2 is the cognate cellular receptor for SARS-CoV and SARS-CoV-2 coronaviruses, and a recently proposed drug target for SARS-CoV-2 (Monteil *et al.*, 2020). Patients presenting with COVID-19 frequently harbor comorbidities such as hypertension, diabetes mellitus, and coronary heart diseases, all which have been associated with increased disease severity (Fang *et al.*, 2020, Ferrario *et al.*, 2005). Since these comorbidities are frequently treated with ACE pathway perturbing medications (ACEIs and ARBs), one possibility is that these medications may make patients more susceptible to SARS-CoV-2 infection.

Though previous epidemiological studies have reported increased mortality and morbidity in COVID-19 patients with hypertension, we sought to examine this risk in the context of ACEI or ARB use (Patel and Verma, 2020). Our retrospective clinical analysis shows that ACEI/ARB exposure increased COVID-19 risk, but showed equivocal effects for morbidity and mortality when conditioning on hypertension status. A recent study reported a protective effect of ACEI/ARB exposure on mortality (Zhang *et al.*, 2020). We were able to replicate their result by conditioning on hypertension. However, this same analysis also revealed no effect, or possibly even a risk, from ACEI/ARB exposure on mechanical respiration and a strong risk of infection. Further, the association with infection was consistent regardless of conditioning on hypertension and in a direct comparison to CCB exposure, mitigating some of the concerns about confounding risk. Taken together, the results from these studies suggest caution should be taken before any clinical guidelines are put in place. Prospective clinical trials or multi-centered blinded studies may be necessary before more concrete conclusions can be drawn. For example, if some patients are more susceptible because they are already expressing high levels of *ACE2*, this could help with targeting ACE2 in these patients as a prophylactic method. However, if the cells respond to infection with *ACE2* expression, and this leads to the cytokine storm seen in patients, then this could be used as a downstream treatment (post-infection), for when ACE2 interacts with TMPRSS2, such as the ongoing trials with camostat mesylate (Hoffman *et al.*, 2020).

Finally, it is notable that the majority of the testing for SARS-CoV-2 so far has relied on nasopharyngeal specimen collection, yet preliminary results here and elsewhere demonstrate SARS-CoV-2 detection from oral specimens is feasible and likely optimal (Woelfel *et al.*, 2020, Wylie et al., 2020). Further studies comparing nasopharyngeal, oropharyngeal, buccal, and saliva collection approaches, as well as a comparison of different swab types, are needed. Depending on the availability of reagents and resources, as well as automation, a LAMP-based approach on such sample types could allow facilities to increase testing capabilities by orders of magnitude. Since viral pandemics can have significant, long-lasting detrimental impacts for affected countries, it is crucial to deploy methods that can track and profile cases (e.g. RNA-seq, LAMP, qRT-PCR) and provide a comprehensive view of host and viral biology. These methods can help mitigate the medical and socioeconomic harm from viral outbreaks, as well as establish protective surveillance networks that can help defend against future pandemics.

## Acknowledgements

We thank the Core Facilities at Weill Cornell Medicine, the Clinical Laboratories at New York Presbyterian Hospital, the Scientific Computing Unit (SCU), OneCodex, the XSEDE Supercomputing Resources and the GISAID Initiative curators and submitters (**Supp Table 10**). We also thank New England Biolabs for providing the reagents for preliminary testing of the LAMP protocols, as well as Eileen Dimalanta and Ted Davis for technical discussions. The authors wish to thank the following members of the HudsonAlpha Discovery team who supported the RNASeq experiments described in the manuscript: Colleen Cowan, John Mote, Arianna Pionzio, Melanie Robinson, and Madison Robison.

## Author Contributions

CEM led the study design and coordination and HR led the clinical collection and validation work in the NYP CLIA laboratory, as well as with MI. Overall supervision and protocol development and implementation for the Zymo RNAClean and NEB assays (SL). DB and CMz performed the LAMP experiments to validate the assay, established a method to quantify LAMP output, and developed a protocol for clinical use of the assay. CW, DX, PR, JG, and JX assisted with sample preparation and sequencing. CeM, DD, JF, AS, JR, MM, EA, IH, DM, MI, BWL, MZ, UG, NPT, NAI, CEM performed analyses. DD, CMz, NAI, MS, BY, KR, CB coordinated and collected environmental samples. EA submitted the IRB application and helped with clinical coordination. Help and insights for analysis from EA, SL, MI, MS, LFW, ML, MC, HR, KR all led to the figures and analyses. All authors reviewed, edited, and approved the manuscript.

## Conflicts of Interest

Nathan Tanner and Bradley W. Langhorst are employees at New England Biolabs.

### IRB

Samples were collected and processed through the Weill Cornell Medicine Institutional Review Board (IRB) Protocol 19-11021069. Observational cohort analysis (ACEI/ARB) was done through the Columbia University IRB Protocol AAAL0601.

### Data Accessibility

Viral sequences were uploaded into GISAID (Global Initiative on Sharing All Influenza Data) site (https://www.gisaid.org), and patient data is being deposited into dbGAP (accession pending).

## Materials and Methods

### Sample Collection and Processing

Patient specimens were collected with patients’ consent at New York Presbyterian Hospital-Weill Cornell Medical Center (NYPH-WCMC) and then processed for qRT-PCR. Nasopharyngeal (NP) swab specimens were collected using the BD Universal Viral Transport Media system (Becton, Dickinson and Company, Franklin Lakes, NJ) from symptomatic patients.

### Extraction of Viral RNA and qRT-PCR detection

Total viral RNA was extracted from deactivated samples using automated nucleic acid extraction on the QIAsymphony and the DSP Virus/Pathogen Mini Kit (QIAGEN). One step reverse transcription to cDNA and real-time PCR (RT-PCR) amplification of viral targets, E (envelope) and S (spike) genes and internal control, was performed using the Rotor-Gene Q thermocyler (QIAGEN).

### Twist Synthetic RNAs

We used two fully synthetic RNAs made by in vitro transcription (IVT) from Twist Biosciences, which were synthesized in 5kb pieces with full viral genome coverage of SARS-CoV-2. They were sequence verified to ensure >99.9% viral genome coverage, and come as 1,000,000 copies per μL, 100μL per tube. The two controls are from Wuhan, China (MN908947.3) and Melbourne, Australia (MT007544.1). Reference sequence designs came from NCBI: https://www.ncbi.nlm.nih.gov/nuccore/MT007544 and https://www.ncbi.nlm.nih.gov/nuccore/MN908947.3.

### Reverse Transcriptase, quantitative real-time PCR (RT-PCR)

Clinical samples were extracted as described above and then tested with qRT-PCR using primers for the E (envelope) gene, which detects all members of the lineage B of the beta-CoVs, including all SARS, SARS-like, and SARS-related viruses, and a second primer set for the S (spike) gene, which specifically detects the SARS-CoV-2 virus. The reaction also contains an internal control that served as an extraction control and a control for PCR inhibition.

Samples were annotated using qRT-PCR cycle threshold (Ct) value for SARS-CoV-2 primers. Subjects with Ct less than or equal to 18 were assigned “high viral load” label, Ct between 18 and 24 were assigned “medium viral load” and Ct between 24 and 40 were assigned “low viral load” classes, with anything above Ct of 40 was classified as negative. We also predicted a combined viral load score using Ct, GloMax QuantiFluor readout from LAMP experiments and fraction of SARS-CoV-2 matching NGS reads in a sample. For this score (40-Ct), (LAMP readout) and (log10(SARS-CoV-2 fraction + 1e-6)) were all normalized between zero and one individually, and summed together using a combination weight of 5 for Ct, 3 for LAMP and 2 for NGS.

### LAMP Primer Sequences

Primers were designed using PrimerExplorer (v4.0), as per guidelines in Zhang *et al.*, 2020. This specifically utilized the LAMP-compatible primers for the on the COVID-19 reference genome (NCBI). LAMP’s inherent specificity (using 4-6 primers vs. 2 for PCR amplification) in combination with this *in-silico* analysis revealed there is limited opportunity for cross-reactivity to allow for false-positive reporting or affect performance of the N-gene primers for SARS-CoV-2 detection (**Supp Table 5**). Overall, the primers had less than 80% homology with the vast majority of tested pathogen sequences. For any organisms where a primer hit >80% homology, only one of the primers (forward or reverse) had significant homology making an amplified product extremely unlikely. Overall, the results of this analysis predict no significant cross-reactivity or microbial interference. We also assessed the potential impact of sequence variation in circulating strains that might lead to poor amplification. In the thousands of sequences deposited in GISAID (Shu and McCauley, 2017), only one site in the priming region was observed to be polymorphic. The polymorphism (T30359C) was only observed in 106 of 6753 (<2%) sequences with coverage of this region. This variant overlaps the priming site of the LB primer but is not near a 3-prime end and is not anticipated to cause amplification failure. The data from Figure 1 show the use of a single-tube, dual-primer protocol, wherein both the N2 gene and E gene primers are present.

#### Primer Sequences: rActin (5′-3′)

ACTB-F3 AGTACCCCATCGAGCACG

ACTB-B3 AGCCTGGATAGCAACGTACA

ACTB-FIP GAGCCACACGCAGCTCATTGTATCACCAACTGGGACGACA

ACTB-BIP CTGAACCCCAAGGCCAACCGGCTGGGGTGTTGAAGGTC

ACTB-LoopF TGTGGTGCCAGATTTTCTCCA

ACTB-LoopB CGAGAAGATGACCCAGATCATGT

#### Gene E Primer Set (5’-3’)

E1-F3 TGAGTACGAACTTATGTACTCAT

E1-B3 TTCAGATTTTTAACACGAGAGT

E1-FIP ACCACGAAAGCAAGAAAAAGAAGTTCGTTTCGGAAGAGACAG

E1-BIP TTGCTAGTTACACTAGCCATCCTTAGGTTTTACAAGACTCACGT

E1-LB GCGCTTCGATTGTGTGCGT

E1-LF CGCTATTAACTATTAACG

#### Gene N2 Primer Set (5’-3’)

N2-F3 ACCAGGAACTAATCAGACAAG

N2-B3 GACTTGATCTTTGAAATTTGGATCT

N2-FIP TTCCGAAGAACGCTGAAGCGGAACTGATTACAAACATTGGCC

N2-BIP CGCATTGGCATGGAAGTCACAATTTGATGGCACCTGTGTA

N2-LF GGGGGCAAATTGTGCAATTTG

N2-LB CTTCGGGAACGTGGTTGACC

### The LAMP Reaction Setup

For each well or Eppendorf tube, we utilized a set of six primers (above) for Gene N, the M1800 LAMP Master Mix (NEB), water, and 11.5μL of the sample. The protocol is as follows:

1. Reagents added:
  a. 12.5 µL M1800 LAMP mix (NEB)
  b. 1-5 µL LAMP primers (Gene N or N2/E mix)
  c. 1-11.5 μL of sample
  d. Remaining volume (to 25 µl) H_2_O
2. Vortex, spin down;
3. Place on Thermocycler at 65°C for 30 minutes with lid at 105 °C;
4. Remove tubes, place on ice for 5 seconds;
5. Visualize over lab bench/ice/paper.

### Oropharyngeal Lysate LAMP Run

Nasopharyngeal and/or oropharyngeal swab samples from 201 patients were collected using a dry cotton swab (cliniswab DS, Aptaca Spa (Italy), #2170/SG/CS). Crude extraction was performed according to pending unpublished European patent [No. 20168 593.0]. In summary, the dry swab was transferred to a 15 ml falcon (Greiner Bio-one, #188.271), filled with 0.5 ml saline solution, and shaken vigorously for 30 min. Afterwards the liquid is transferred to a screw-cap (Sarstedt, #72.692.005). 10µl of the crude RNA extract was added to 12.5 µl 2x NEB LAMP master mix (#M1800L), 2.5 µl of water, 1 µl 25x primer master mix Gen N, and was incubated at 65 °C for 30-40 min. A sample of a patient whom tested positive using an approved qRT-PCR test (sample #1123) was used as an internal control. The read-out was performed visually by color change from pink to yellow or orange.

The RNA isolation was performed with 50 µl of the crude extract on the QIAsymphony with the DSP Virus/Pathogen Kit (Qiagen, #937055). The RT-PCR was performed using 5 µl of 85 µl eluate with TIB MolBiol Lightmix® MODULAR SARS AND WUHAN CoV E-Gene Kit. Analysis was done with the LightCycler(R) 480 II software and calculated CP values were used for statics and graphical analysis. The Standard curve with a synthetic RNA control (Twist Bioscience, #MT007544.1) was generated using the lamp assay and in parallel with the qRT-PCR. The control RNA was diluted serially 10-fold with water and the absolute copy number, ranging from 10^5^ to 10^−5^ was analyzed. Gel electrophoresis after visual read-out of the LAMP assay was done by loading 5 µl of the lamp reaction with 5 µl 2x loading dye on a 1.5 % agarose (Seakem LE Agarose, Lonza #50004) together with 5 µl of a 1kB DNA Ladder (Roche). The electrophoresis was performed at 90 V constant.

The photometric read-out of the standard curve was performed in a plate reader. To this end the lamp reaction was transferred into a 96-well V-shaped cell-culture plate (Greiner Bio-One, #651180). After measuring the absorbances at 432nm and 560nm, the ratio abs(432)/abs(560) was calculated. The relative absorbance abs(432)/abs(560) was calculated by subtraction of the negative (water) control from all samples (including the negative control). All values above a threshold of 0.1 are considered as a positive assay read-out and are marked with “+”, all other values are negative and are marked “-”. Statistical and graphical analysis were performed with GraphPad Prism 8.0.4.

### Light Intensity and Data Processing

Completed reactions were analyzed with the Promega GloMax Explorer (Promega GM3500) fluorometer using the QuantiFluor ONE dsDNA system (Promega E4871). This system recorded fluorometric readout from each well using an emission filter of 500-550nm, an excitation filter set at blue 475 nm, and a high sensitivity setting on the Glomax software. Values were then tabulated and compared with controls (positive and negative). The intensity threshold of 2.5x negative control was used as the threshold for positive detection.

### DNAse treatment, rRNA depletion, and RNAseq library construction

For library preparation, all samples’ total nucleic acid (TNA) were treated with DNAse 1(Zymo Research, Catalog # E1010), which cuts both double-stranded and single-stranded DNA. Post-DNAse digested samples were then put into the NEBNext rRNA depletion v2 (Human/Mouse/Rat), Ultra II Directional RNA (10ng), and Unique Dual Index Primer Pairs were used following the vendor protocols from New England Biolabs (except for the first flowcell, see supplemental figures). Kits were supplied from a single manufacturer lot. Completed libraries were quantified by Qubit or equivalent and run on a Bioanalyzer or equivalent for size determination. Libraries were pooled and sent to the WCM Genomics Core or HudsonAlpha for final quantification by Qubit fluorometer (ThermoFisher Scientific), TapeStation 2200 (Agilent), and QRT-PCR using the Kapa Biosystems Illumina library quantification kit.

### Taxonomic Classification of Sequence Data

All complete genome or chromosome level assemblies from RefSeq database for archaea, bacteria, protozoa, fungi, human and viruses including SARS-CoV and SARS-CoV-2 genomes were downloaded and used for building a classification database for Kraken2 (k=35, ℓ=31) (O’Leary *et al.*, 2016; Wood *et al.*, 2019).

To get an approximation for the positive and negative classification rate, the BBMap random-reads script was used to simulate 10 million 150bp paired-end Illumina reads from the database sequences (Segata *et al.*, 2016). For the negative test all sequences in the database excluding SARS-CoV and SARS-CoV-2 genome were removed from the sequences and the simulated reads were mapped with the Kraken2 database (Supp Table 1).

For the positive test, the same process was repeated using only SARS-CoV-2 genome (**Supp. Table 2**). Positive results show >99% of SARS-CoV-2 reads uniquely map to either SARS-CoV or SARS-CoV-2, with the remaining 1% are ambiguous, potentially matching multiple taxa (**Supp. Table 2**). All sequences were classified using the Kraken2 database. To remove the potential contamination of reads that are homologous across multiple species we used Kraken2 outputs to filter sequences to either human (uniquely matching Homo sapiens and no other taxon in our database), SARS-CoV-2 (either matching SARS-CoV or SARS-CoV-2 due to homology between these two viruses), and remaining reads that may be coming from unclassified, archaeal, bacterial, viral, fungal, protozoan or ambiguously mapping reads to human or SARS-CoV (*Li*, 2015).

Using kraken2 classifications common respiratory pathogens were identified in clinical samples. Any SARS-CoV-2 negative sample with >0.01% relative abundance (normalized after the exclusion of any human, SARS-CoV-2 and uncharacterized reads) for presence of viral pathogens were classified as potential unrelated viral infection (**Supp. Figure 7**). These samples were used as controls during specific differential expression comparisons to compare the common effects of viral infections on host cells from SARS-CoV-2 infection.

### Viral Genome Assembly

Reads unambiguously mapping to SARS-CoV or SARS-CoV-2 were aligned to the Wuhan-Hu-1 (Genbank accession MN908947.3) reference using bwa mem (Li, 2013). Variants were called using iVar, and pileups and consensus sequences were generated using samtools (Li *et al.*, 2009; Grubaugh *et al.*, 2019; Greenfield *et al.*, 2020). Any sample with >99% coverage above 10x depth for SARS-CoV-2 genome were taken as reliable samples, which resulted in 155 samples (146 positive, 9 negative). 155 clinical samples were compared to 7,797 SARS-CoV-2 sequences from GISAID (as of April 23, 2020) (9, 10). All sequence filtering, alignments, phylogenetic inference, temporal ordering of sequences and geographic reconstruction of likely transmission events were done using Nextstrain (Katoh and Standley, 2013; Sagulenko *et al.*, 2018; Hadfield *et al.*, 2018). Sequence identity and coverage metrics were calculated using Mview (Brown *et al.*, 1998). Phylogenetic trees were created using Nextstrain’s augur as described above, and visualized using the ggtree package in R (Yu, 2020).

### Viral variant calling and allelic fraction estimation

Full-length viral consensus sequences were aligned to the Wuhan-Hu-1 reference using bwa mem (Li, 2013) with default settings. Variants were identified by enumerating the coordinates and query / reference subsequences associated with mismatches (SNV) and gaps in the query (deletion) and reference (insertions) using R/Bioconductor (GenomicRanges, Rsamtools, Biostrings packages) and Imielinski lab gChain packages (https://github.com/mskilab/gChain). Exhaustive variant calling on read alignments was additionally performed using bcftools mpileup and call, with variant read support (VAF, alternate allele count) enumerated with the R/Bioconductor Rsamtools package.

For each variant, a posterior distribution of variant allele fraction (VAF) was computed using a beta distribution with shape parameters *θ* comprising reference and alternate allele counts and pseudo count of 0.5. Variants were classified as het (heterogenous) if *P*(*VAF* > 0.05 *∧ VAF* < 0.95 | *θ*) > 0.95. For a given specimen, posterior VAF distributions of *k* heterogenous variants were then integrated using a histogram density estimator by summing the posterior VAF density across all variants at each point of a grid of 100 points evenly spaced in the (0,1) line. This (unnormalized) mixture density was visualized alongside the individual VAF densities as an estimate of the probability density of a putative viral subclone.

### SARS-CoV-2 genome wide association study

Population reference and alternate allele counts were computed for all SNV and indel variant calls detected in at least three cases in the A2 clade subset of our NYPH-WCMC sequences (123 cases, see above). Similar counts were computed across 537 New York and 7269 non-New York sequences that were obtained from GISAID and incorporated into the Nextstrain tree (see above). A one-tailed Fisher’s Exact test was used to compare allele counts in NYPH-WCMC vs. non-New York GISAID cases. All variants with FDR < 0.1 were reported as significant. Significant variants from the initial analysis were replicated in a similar comparison of New York vs non-New York GISAID sequences.

### Cell Deconvolution Analysis

Bulk RNAseq count data was deconvoluted into cell composition matrices using the MUSIC algorithm (Wang *et al.*, 2019) on a reference single cell RNAseq data from upper respiratory epithelium obtained from nasal brushes and upper airway and lung cells ((Vieira *et al.*, 2019; Lukassen *et al*. 2020).

### Human Transcriptome Analysis

The reads that mapped unambiguously to the human reference genome via Kraken2 were used to detect the host transcriptional response to the virus. Reads matching *Homo sapiens* were trimmed with TrimGalore, aligned with STAR (v2.6.1d) to the human reference build GRCh38 and the GENCODE v33 transcriptome reference, gene expression was quantified using featureCounts, stringTie and salmon using the nf-core RNAseq pipeline (Pertea *et al.*, 2015; Malinen *et al.*, 2005; Johnson *et al.*, 2007; Robinson *et al.*, 2010; Naccache *et al.*, 2014; Zamani *et al.*, 2017; Ewels *et al.*, 2019). Sample QC was reported using fastqc, RSeQC, qualimap, dupradar, Preseq and MultiQC (Okonechnikov *et al.*, 2016; Andrews, 2015; Ewesl *et al.*, 2016; Sayols *et al.*, 2016; Wang *et al.*, 2012). Samples that had more than 10 million human mapped reads were used for differential expression analysis. Reads, as reported by featureCounts, were normalized using variance-stabilizing transform (vst) in DESeq2 package in R for visualization purposes in log-scale (Love *et al.*, 2014). Limma voom and DESeq2 were used to call differential expression with either Positive cases vs Negative, or viral load (High/Medium/Low/None excluding any samples with evidence of other viral infections) as reported by either qRT-PCR cycle threshold (Ct) values, or using the inverted normalized Ct value as continuous response for viral levels, where Ct of 15 is 1.0 and Ct of >40 is taken as 0 (Law *et al.*, 2014). Genes with BH-adjusted p-value < 0.01 and absolute log2 fold-change greater than 0.58 (at least 50% change in either direction) were taken as significantly differentially regulated (Benjamini and Hochberg, 1995). The same approaches were repeated correcting for potential confounders in our data in two ways. In the first correction ciliated cell fraction (as predicted by MUSIC) was added as another covariate to our model. For the second correction SVA was run on the data and the resulting two surrogate variables were included in a multivariate model (Leek *et al.*, 2012). The complete gene list for all comparisons are given in Supp Table 3. Resulting gene sets were ranked using log2 fold-change values within each comparison and put into GSEA to calculate gene set enrichment for molecular signatures database (MSigDB), MGI Mammalian Phenotypes database and ENCODE transcription factor binding sets (Liberzon *et al.*, 2011; Subramanian *et al.*, 2005; Kuleshov *et al*. 2016; Sergushichev, 2016; Smith *et al.*, 2018). Any signature with adjusted p-value < 0.01 and absolute normalized enrichment score (NES) >= 1.5 were reported (Supp Table 3).

### Cross-reactivity Analysis

Primers were compared with a list of sequences from organism from the same genetic family as SARS-CoV-2 and other high-priority organisms listed in the United States Food and Drug Administration’s Emergency Use Authorization Template (https://www.fda.gov/media/135900/download). Using the sequence names in the EUA template, the NCBI taxonomy database was queried to find the highest quality representative sequences for more detailed analysis. Primers were compared to this database using Blast 2.8.1 and the following parameters (word size: 7, match score: 2, mismatch score: −3, gap open cost: 5, gap extend cost: 2). Up to 1000 hits with e-value > 10 were reported.

### Inclusivity Analysis

Unique, full-length, human-sample sequences were downloaded from the GISAID web interface. These sequences were aligned to NC_045512.2 (Wuhan SARS-CoV-2) using minimap2 -x asm5 and visually inspected using IGV 2.8.0 with allele frequency threshold set to 0.01.

### ACE Inhibitor/Angiotensin Receptor Blocker Cohort Analysis

We compared usage of ACE inhibitors (ACEI) and angiotensin receptor blockers (ARB) in an observational cohort analysis of 8,856 patients with suspected SARS-CoV-2 infection (4,829 of which tested positive). ACEIs and ARBs are commonly taken continuously for several years (Bonarjee et al., 2001). We defined a cohort of ACEI/ARB exposed patients as those that have an ACEI/ARB prescription order sometime after January 1st, 2019. We compared the frequency of ACEI/ARB exposure in three cohort comparisons:

i. SARS-CoV-2 tested positive patients versus SARS-CoV-2 tested negative patients,
ii. SARS-CoV-2 positive patients who require mechanical ventilation versus those who did not, and
iii. SARS-CoV-2 positive patient survival versus death.

In addition, we perform one post-hoc comparison to evaluate the individual effects of particular ACEIs or ARBs among SARS-CoV-2 positive patients who are exposed.

### Cohort and Data Source

Our cohort data for SARS-CoV-2 suspected patients is extracted from the electronic health records at NYPH-CUIMC. We used data collected starting on March 10th, 2020 through April 20th, 2020. In addition, we used data from 90,989 patients, who were not tested for SARS-CoV-2 infection and were 18 years or older, with available electronic health records from January 1st, 2019 through September 24th, 2019 to represent a comparison population of patients. In both cases, data extracted included disease diagnoses, laboratory measurements, medication and pharmacy orders, and patient demographics. We derived mortality from a death note filed by a resident or primary provider that records the date and time of death. Intubation was used as an intermediary endpoint and is a proxy for a patient requiring mechanical respiration. We used note types that were developed for patients with SARS-CoV-2 infection to record that this procedure was completed. We validated outcome data derived from notes against the patient’s medical record using manual review.

### Experimental Statistical Methods

We conducted univariate analysis of the frequency differences of ACEI/ARB exposure and multivariate regression analysis to account for established risk factors for SARS-CoV-2 outcomes (i.e. age, sex, baseline IL-6 upon admission, and comorbidities) (Zhou et al., 2020). We use logistic regression for analysis for comparing cohort (i) and performed survival analysis using a Cox proportional hazards model for cohort comparisons (ii) and (iii). We cannot determine from our data the date of infection. For the study start date for the patient, we use the date of testing positive or the first date a diagnosed patient showed symptoms, whichever came first, minus seven days. Quantitative variables (i.e., age and the first IL-6 measurement) are linearly scaled to [0,1] to facilitate comparison of model coefficients to those of dichotomous variables. Prior to conducting our multivariate analysis, we evaluated and removed correlated covariates. Finally, we repeated the previously described analysis for a cohort restricted to patients with a documented history of clinically diagnosed hypertension. We include all model results in the supplemental tables.

### Comorbidity Definitions

Risk factors were assigned using OMOP CDM concept IDs 316866 for “Hypertensive Disorder”, 317576 for “Coronary Arteriosclerosis”, 201820 for “Diabetes mellitus”, and 437525 for “Overweight”. In each case, the concept ID and all descendant concepts were used to define the risk factor phenotype, and individuals were assigned the phenotype if they were assigned any of the codes.

### Statistical and visualization software

All electronic health data analyses were performed in Python 3.7 and all models were fit using R 3.6.3. Survival analyses (Cox regressions and survival curves) were performed with the survival package for R, version 3.1-12. Additional statistical analyses, processing, transformation, and visualization of genomic data were completed in R / Bioconductor (‘Rsamtools’, ‘GenomicRanges’, ‘Biostrings’) and additional Imielinski Lab R packages (‘gTrack’, ‘gChain’, ‘gUtils’, ‘RSeqLib’) available at https://github.com/mskilab. Visualization of phylogenies was completed using Auspice and the ‘ggtree’ and ‘ape’ libraries for R.

## Supplemental Figures and Tables

**Supplemental Figure 1.**
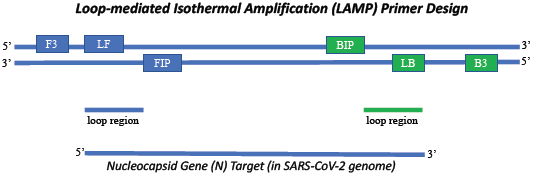
LAMP Primer Design. PrimerDesigner (v4) was used to create a set of six primers that would specifically target the nucleocapsid gene (N) in the SARS-CoV-2 genome. Primer sequences are listed in the methods section.

**Supplemental Figure 2.**
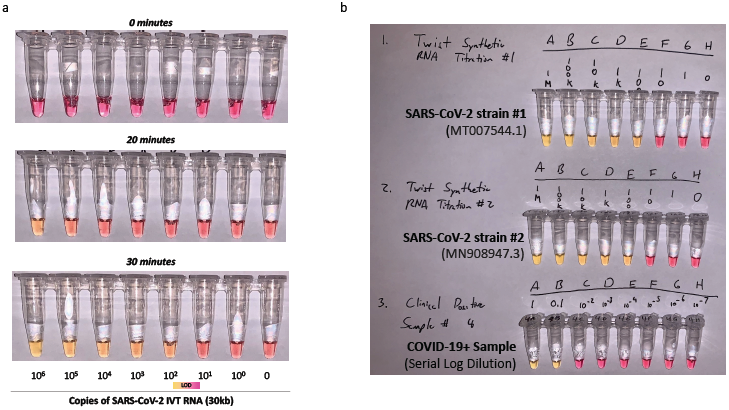
Additional Testing and Titration of the LAMP Assay with Synthetic and Clinical Samples. Samples were prepared using the LAMP protocol with a reaction time of 30 minutes. Reaction progress was measured (a) from 0, 20, and 30 minutes. (b) This was repeated for both Twist SARS-CoV-2 synthetic RNAs (MT007544.1, top and MN908947.3, middle) from 1 million molecules of virus (10^6^), then titrated down by log10 dilutions. Limit of Detection (LOD) range is shown with a gradient after 30 minutes between 10 and 100 viral copies. (bottom) A clinically positive sample that was not detectible by Qubit (<0.05 ng/mL) was nonetheless detected by LAMP, in accordance with detection of low viral titer samples.

**Supplemental Figure 3.**
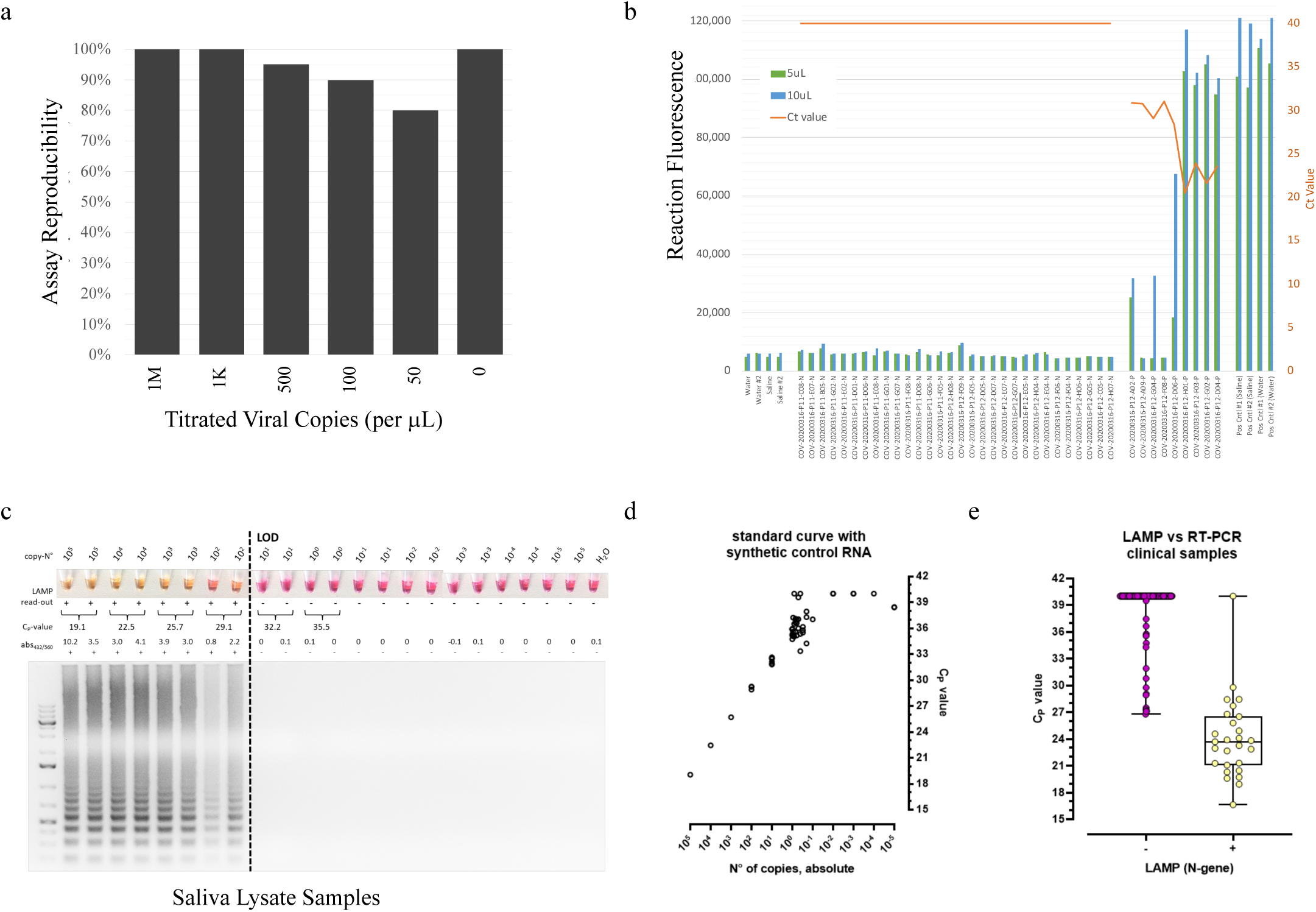
Reproducibility, sensitivity, and specificity for the LAMP Assay. (a) Testing with a synthetic SARS-CoV-2 RNA that was serially titrated and measured in replicates (n=10) showed 100% and 95% reproducibility at 1,000 and 500 copies, respectively, with lower rates at lower viral titers. (b) Replicates of a clinically positive (by qRT-PCR) sample at 10 uL (blue) compared to 5uL (green) showed high concordance, with greater sensitivity with increased reaction volume. (c) Whole oropharyngeal swab lysates from clinical positive (Cp-value >0) and negative samples (Cp = NA) were used to test the LAMP reaction. (d) The standard curve with synthetic RNA was also tested relative to absolute number of copies (x-axis) and the Cp value (y-axis). (e) The Cp value for the LAMP positive (+, right) and negative (-, left) were compared to the Cp value from qRT-PCR (y-axis).

**Supplemental Figure 4.**
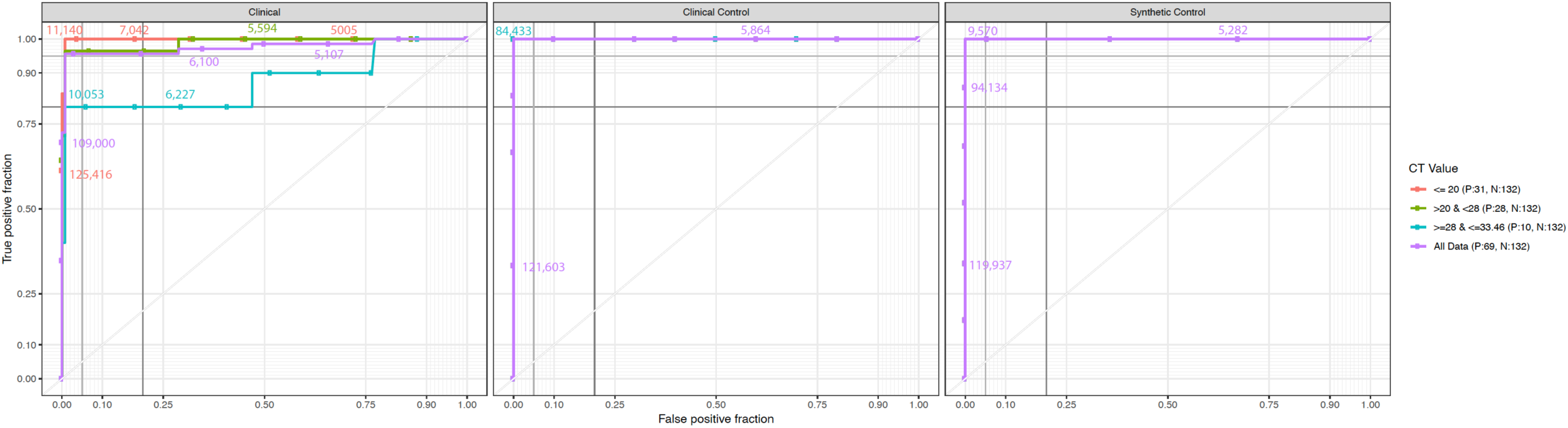
Ct and RFU (Relative Fluorescence Units) Thresholds for the LAMP Assay. With increasing viral load, as measured by qRT-PCR, the LAMP assay shows an increased sensitivity (y-axis) and specificity (y-axis). For high and medium viral count samples (Ct < 28, red and green line), we see 100% sensitivity, and for low viral count, we see 81% sensitivity and 100% specificity (blue). Across all data (purple), we see 95.6% sensitivity and 99.4% specificity. RFU thresholds are shown as numbers.

**Supplementary Figure 5.**
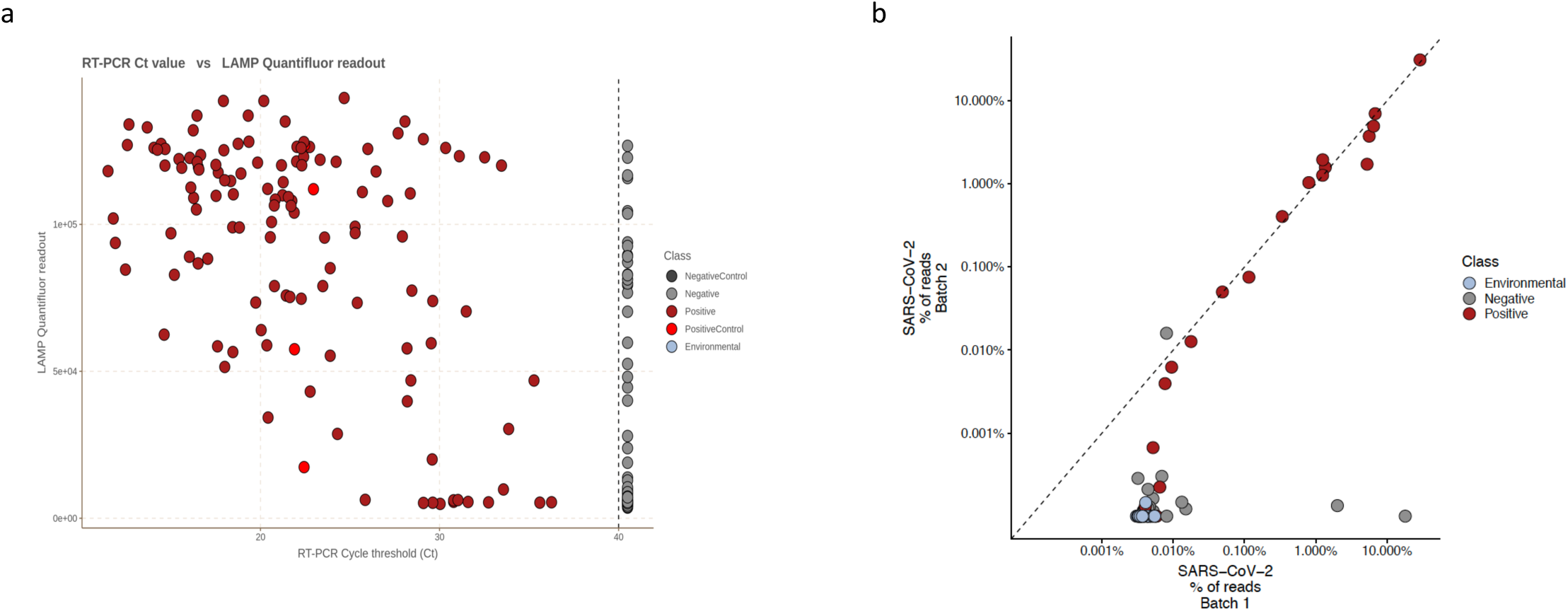
Correlation between LAMP Reaction Output and qRT-PCR as well as RNA-seq replicates. (a) Clinical samples tested by qRT-PCR (Positive, dark red or Negative, light grey) were run with the LAMP assay and compared to the buffer blanks (Negative Control, dark grey), Synthetic RNAs or Vero 6 cell extracts with SARS-CoV-2 infection (Positive Controls, light red), and Subway Samples (Environmental, blue, lower right). The DNA abundance, as measured with the GloMax QuantiFluor (y-axis) is compared to the Ct Threshold for qRT-PCR (x-axis), with lower Ct values representing higher viral abundance. (b) RNA-seq replicates from batch 1 (single index) vs. batch 2 (dual index adapters), showing the broader dynamic range of dual-index libraries (R=0.85).

**Supplementary Figure 6.**
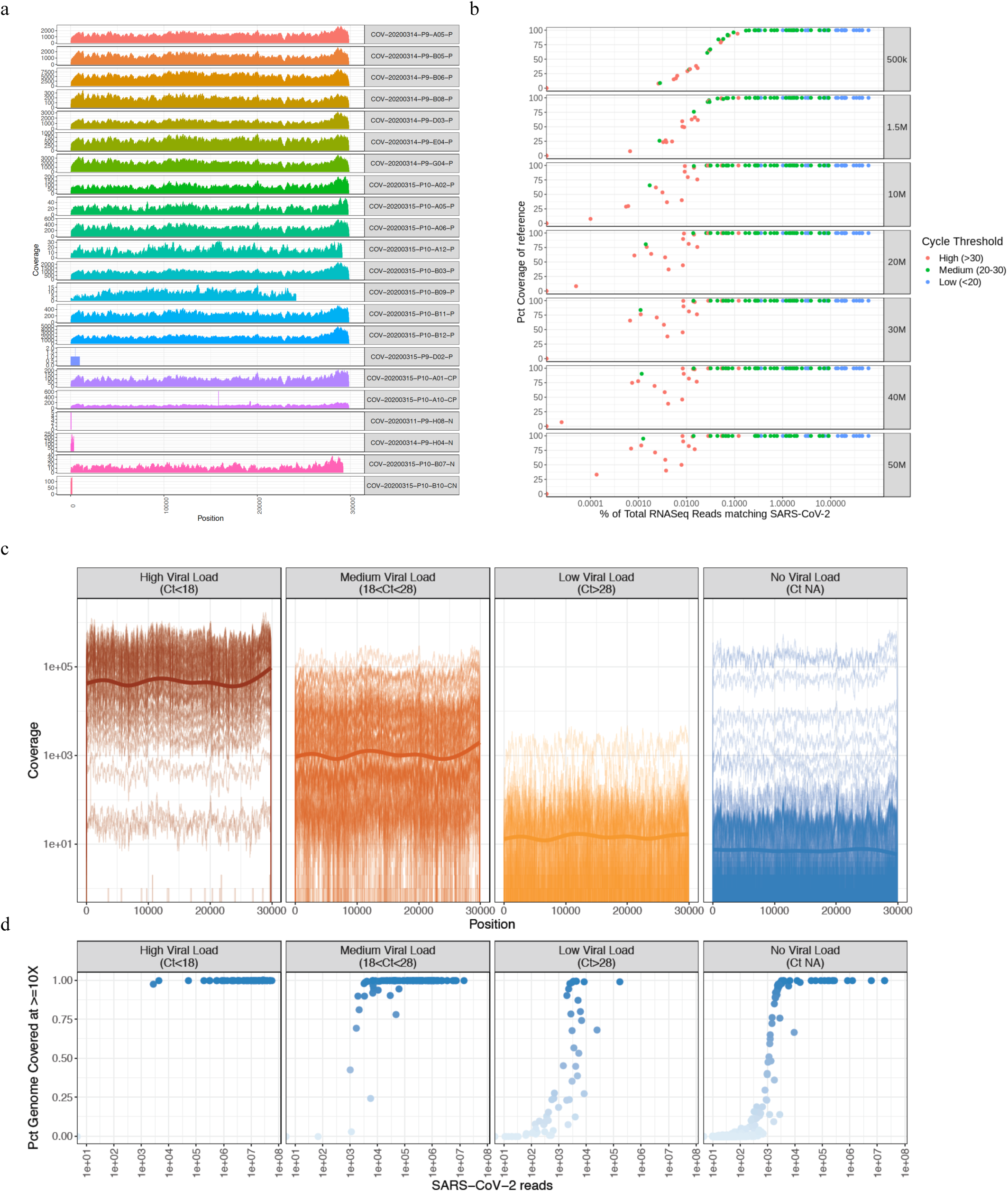
Viral genomes from RNA-seq data and titration of coverage. (a) The coverage plot across the SARS-CoV-2 genomes (viral coordinate on bottom, colored by sample) from a representative set of clinical positive samples. Sample names with the suffixes CN and N are clinical negative (buffer), P are qRT-PCR positive, and CP are Vero E6 cells with virus. (b) Downsampling (right annotation) of the samples and mapping to the SARS-CoV-2 genome to gauge the percent coverage (y-axis) as a function of the viral quantification by qRT-PCR (Ct thresholds, low <20, medium 20-30, and high>30). (c) Average coverage statistics for the low, medium, and high Ct samples, as well as the mean coverage for each of these samples. (d) The cycle threshold (x-axis) vs. the coverage of the genome (y-axis and color depth) for the total RNA-seq.

**Supplemental Figure 7.**
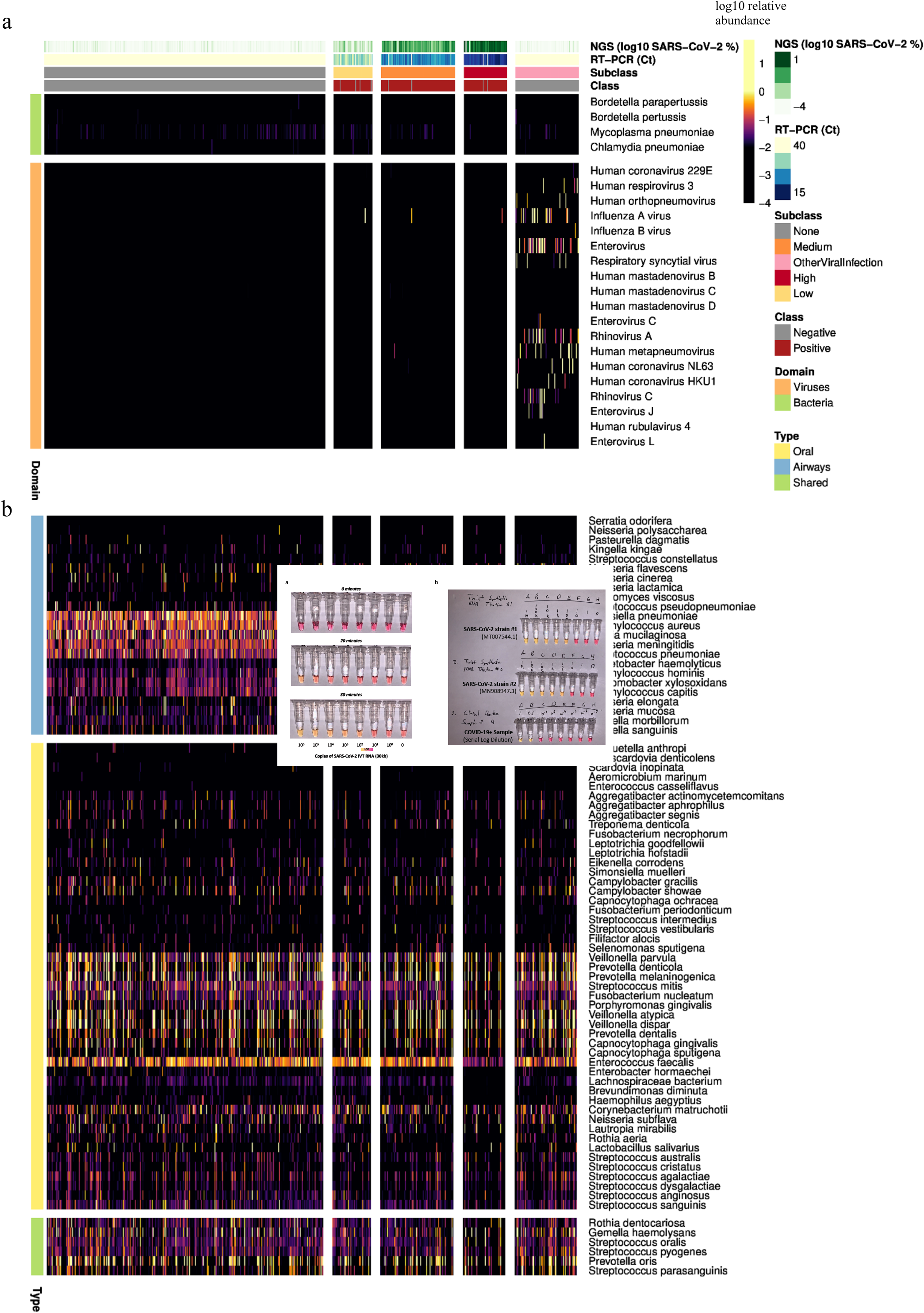
Metatranscriptome profiles of the patient cohorts. (top row, all panels) Samples were quantified by a range of viral detection methods, including RNA-seq (log10 SARS-CoV-2 % of reads) and qRT-PCR (Ct values) to create a three-tier range of viral load for the positive samples (right) compared to the clinically-annotated negative samples (class, red or grey). (bottom) The detected microbial species (horizontal lines) are plotted as log10 abundance. (a) Common respiratory pathogens plotted as a log10 abundance of mapped reads, with each organism as a line and each vertical column as a patient. (b) The same layout of samples as (a), but with bacteria annotated from the Human Microbiome Project (HMP) as normal airway (blue, top portion), oral (yellow, middle set), or both oral and airway flora (green, bottom).

**Supplemental Figure 8.**
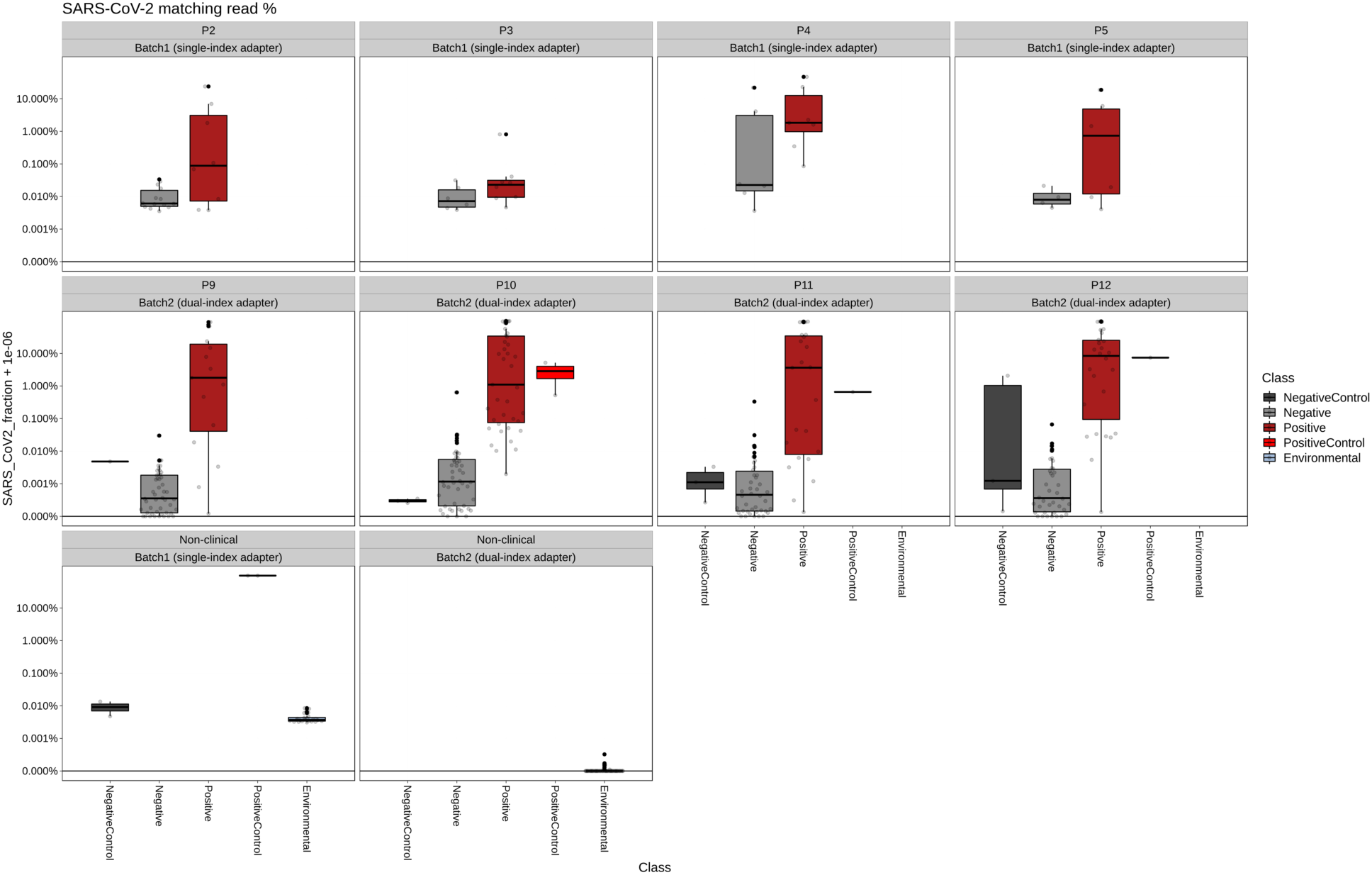
Proportion of RNA-seq reads mapping SARS-CoV-2. Clinical samples tested by qRT-PCR (Positive, dark red or Negative, light grey) were sequenced and compared to the buffer blanks (Negative Control), dark grey, Synthetic RNAs or Vero 6 cell extracts with SARS-CoV-2 infection (Positive Controls, light red), and Subway Samples (Environmental, blue). Read totals are shown on the y-axis. Differences between single index barcodes are plotted across each of the plates of samples that were processed.

**Supplemental Figure 9.**
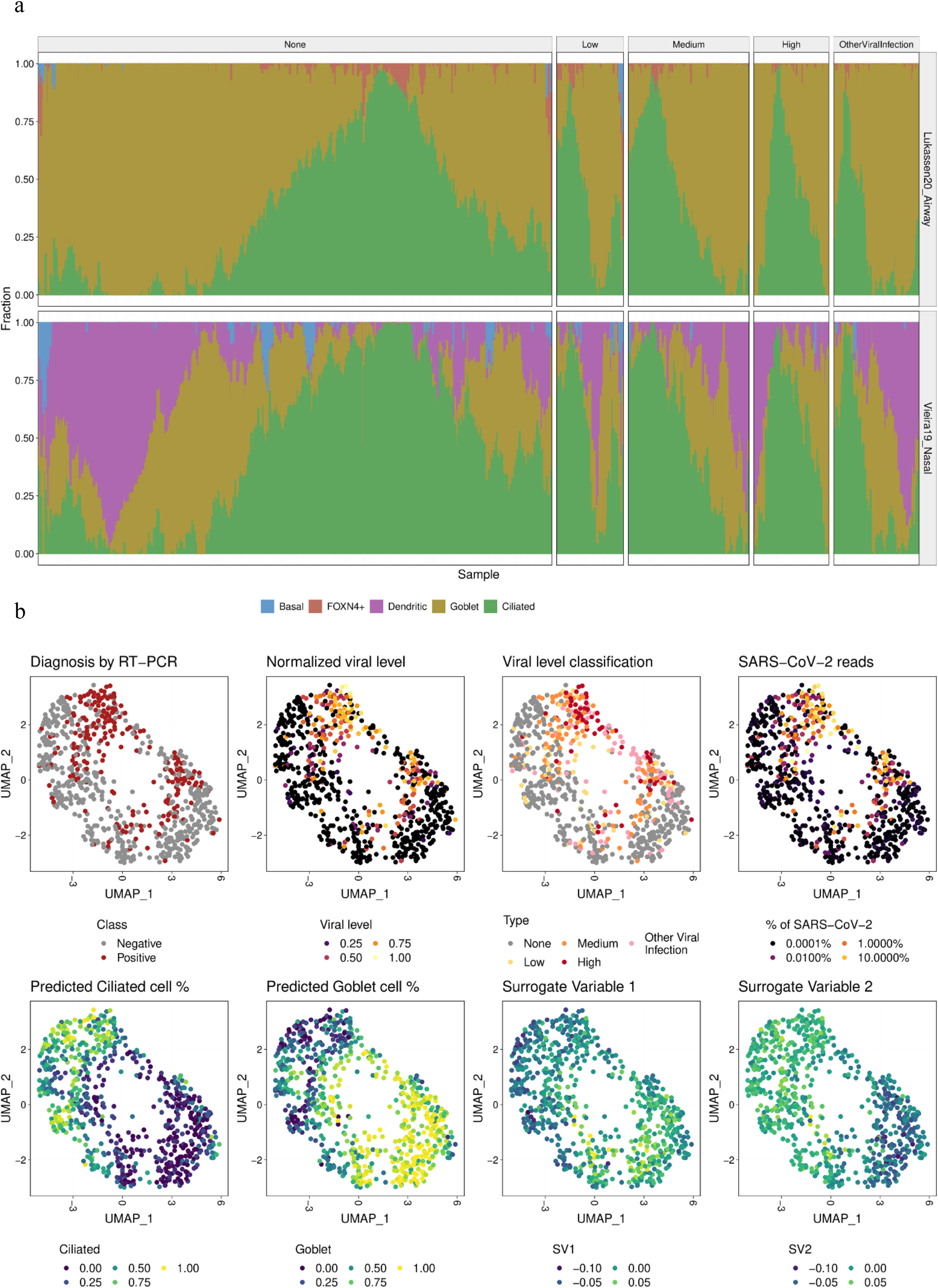
Cellular sub-type deconvolution from nasopharyngeal (NP) swabs. The MUSIC algorithm was used to separate the cellular sub-type gene expression signatures present in the total RNA-seq data from the NP swabs. (a) The fraction of cells estimated for each cell type (y-axis) was calculated for the clinical samples that tested positive by RT-qPCR in different viral levels, as well as negative by RT-qPCR, with proportions shown for goblet cells (yellow), ciliated cells (green), basal cells (blue), dendritic cells (purple) and FOXN4+ cells (red). First row shows deconvolution using the single cell transcription reference set of upper airway cells by Lukassen *et al* 2020. and the second row shows a similar approach using nasal epithelial cells by Vieria *et al*. (b) Clinical samples are embedded in two-dimensions using dimensionality reduction by UMAP and separate features of samples are projected onto the points including RT-PCR based diagnosis, viral levels, fraction of SARS-CoV-2 reads as captured by NGS, predicted ciliated and goblet cell fractions from cell deconvolution using Lukassen airway reference, and the two predicted surrogate variables.

**Supplemental Figure 10.**
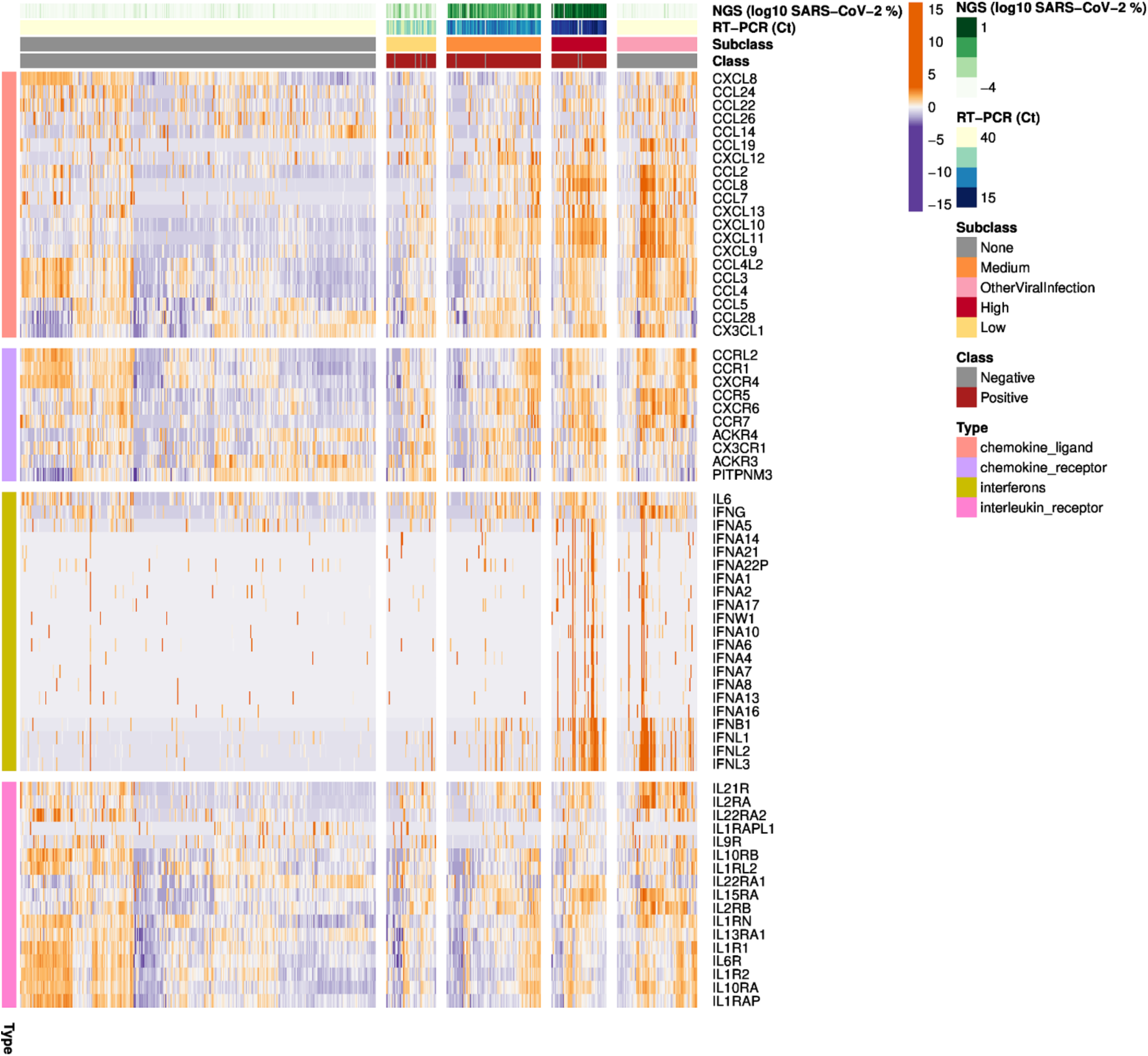
Cytokine and interferon profiles of the host transcriptome. Top rows: samples were quantified by a range of viral detection methods, including LAMP (QuantiFluor), RNA-seq (log10 SARS-CoV-2 % of reads), and qRT-PCR (Ct values) to create a three-tier range of viral load for the positive samples (right) compared to the clinically-annotated negative samples (class, red or grey). (bottom) The differentially expressed genes of SARS-CoV-2 positive patients compared to SARS-CoV-2 negative patients showed up-regulated (orange) genes as well as down-regulated (purple) genes. Heatmap is separated into Chemokine ligand, chemokine receptor, interferon and interleukin receptor profiles for the samples (x-axis) is plotted for each related gene (y-axis).

**Supplemental Figure 11.**
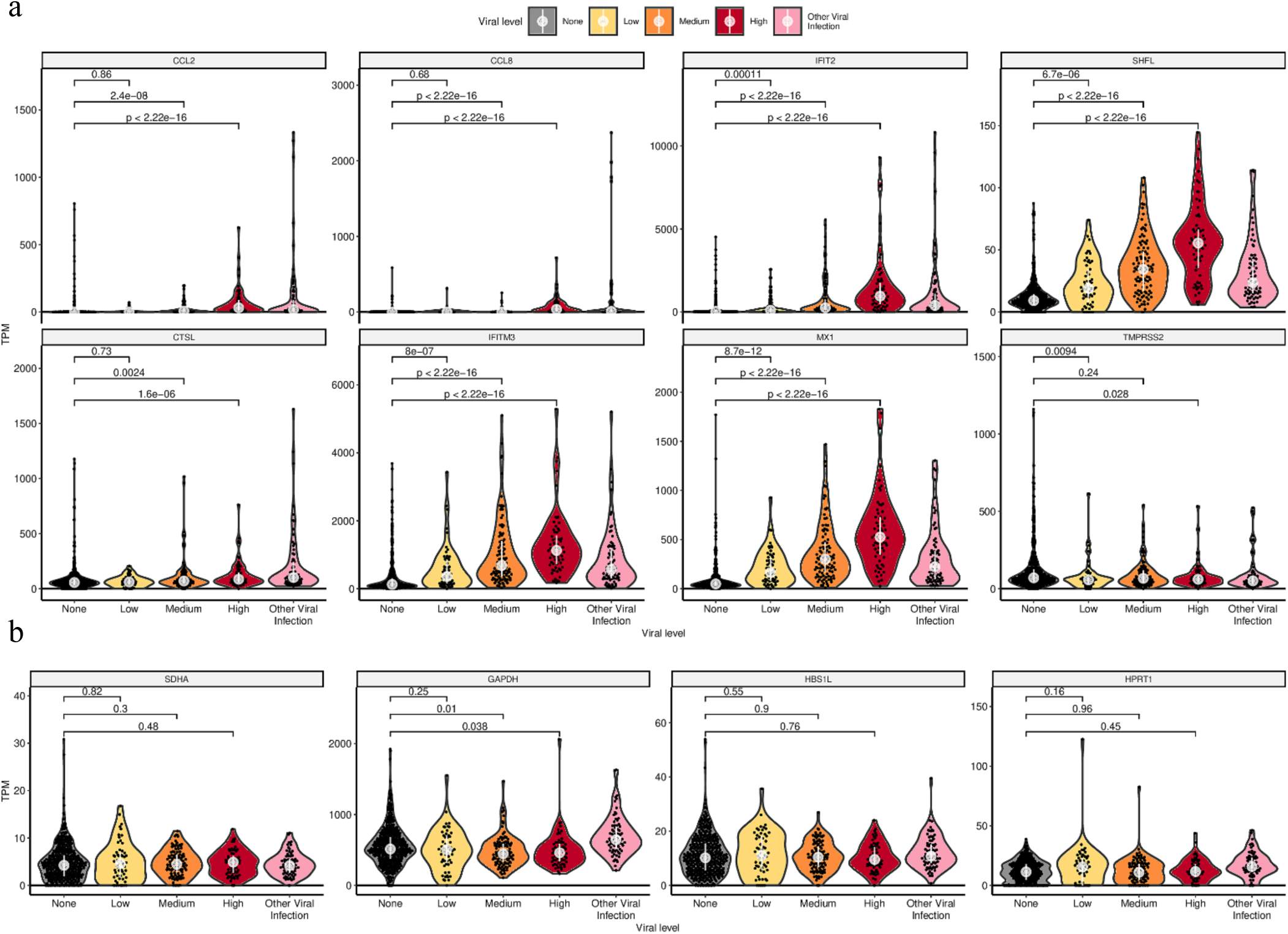
Host gene expression relative to SARS-CoV-2 infection. Clinical samples were sequenced with RNA-seq and quantified to a set of genes for their expression levels. Samples with no virus (grey) were compared to those with low (yellow), medium (orange), and high (red) expression levels, based on qRT-PCR. p-values are calculated by Wilcoxon rank sum test and are not adjusted for multiple testing for a given gene. (a) Additional genes that were differentially expressed, or reported to be important for SARS-CoV-2 entry. (b) Expression of housekeeping genes in different groups.

**Supplemental Figure 12.**
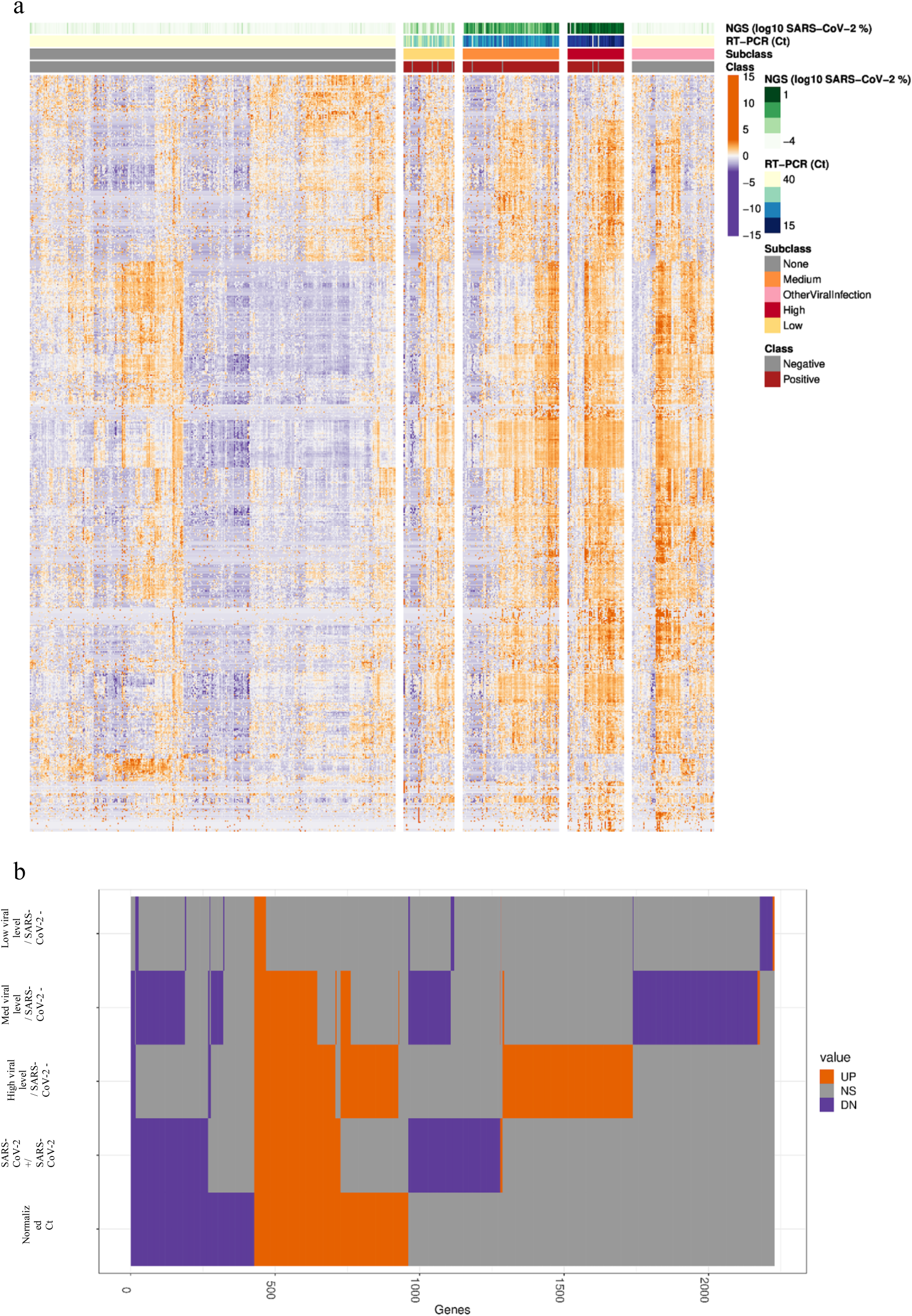
Differentially expressed genes. (a) Top rows: samples were quantified by a range of viral detection methods, including LAMP (QuantiFluor), RNA-seq (log10 SARS-CoV-2 % of reads), and qRT-PCR (Ct values) to create a three-tier range of viral load for the positive samples (right) compared to the clinically-annotated negative samples (class, red or grey). (bottom) The differentially expressed genes of SARS-CoV-2 positive patients compared to SARS-CoV-2 negative patients showed up-regulated (orange) genes as well as down-regulated (purple) genes. All differentially expressed genes with adjusted p-value < 0.001 and |log2 fold-change| > 1.5 (> 2.82 fold) are shown across different patient groups by viral level and presence of other viral pathogens. (b) Intersection heatmap of differentially expressed genes across different comparisons with genes in x-axis and comparisons in y-axis, with a core set of up-regulated genes (orange) distinct form the set of down-regulated genes (purple), compared to genes that are not significantly differently expressed (grey) in any comparison (Limma voom, q-value <0.01, |log2FC| >0.58).

**Supplemental Figure 13.**
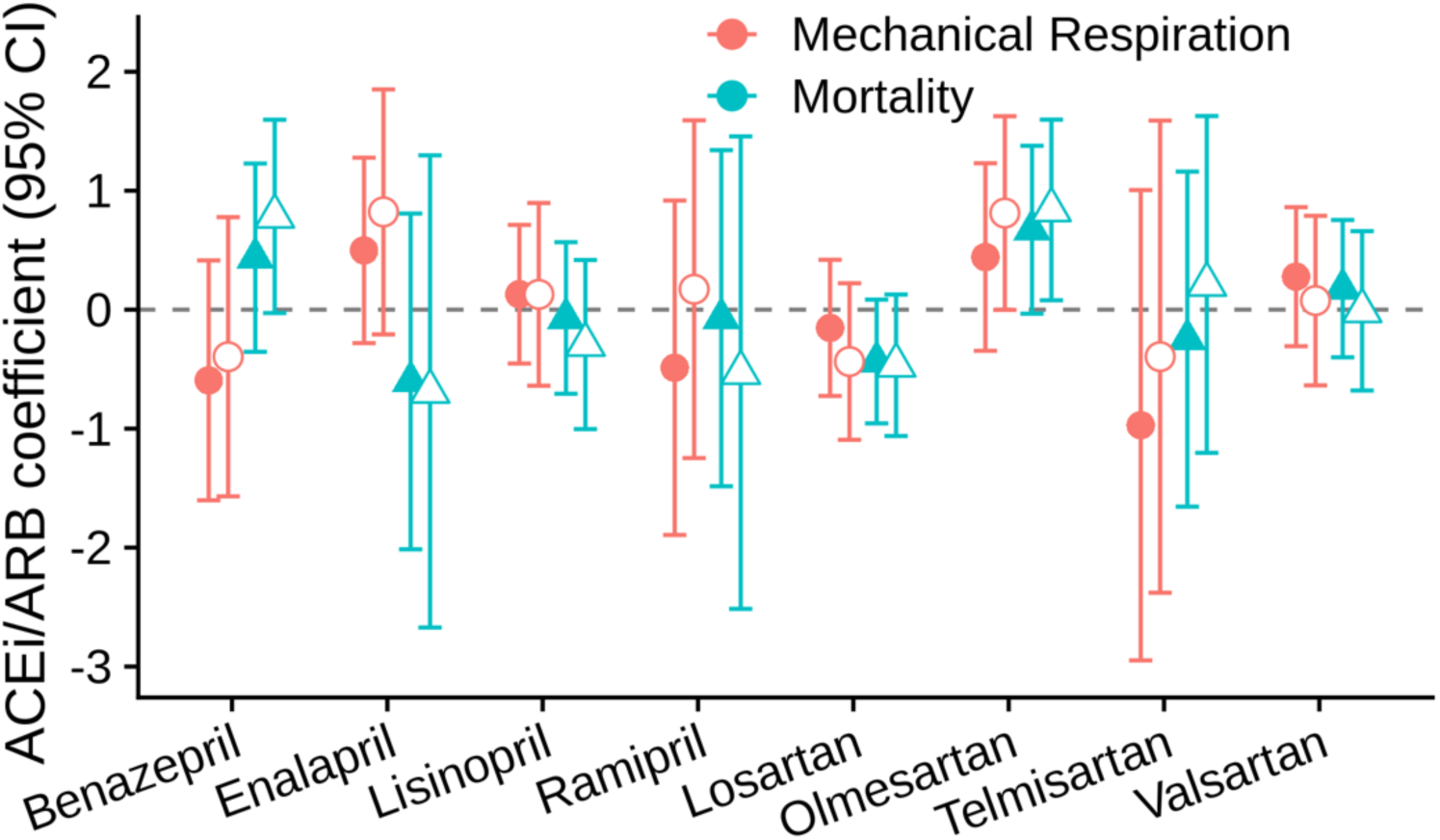
ACEI/ARBs Multivariate and Comparative Analyses. Comparison of the effects of different ACEIs (benazepril, enalapril, lisinopril, ramipril) and ARBs (losartan, olmesartan, telmisartan, valsartan) when compared to patients exposed to other ACEIs or ARBs, respectively. There were no significant findings amongst the different ACEIs. However, among the ARBs we found that olmesartan was associated with a significant increase in mortality (HR=2.3 CI: 1.1-4.9, p=3.05E-02).

**Supplementary Table 1. Metatranscriptome Profiles of All Samples**.

Appended.

**Supplementary Table 2. Taxonomic Mis-assignment Filter**.

Appended.

**Supplementary Table 3. Differentially Expressed Genes in SARS-CoV-2 +/-patients**. Appended.

**Supplemental Table 4. Gene Ontology Pathways**.

Appended.

**Supplementary Table 5. SARS-CoV-2 Rapid Colorimetric LAMP Detection Test: N Primers Specificity**

Appended.

**Supplementary Table 6. Logistic Regression Coefficient**

Appended.

**Supplementary Table 7. Hypertension Logistic Regression Coefficient**

Appended.

**Supplementary Table 8. COX Regression Coefficients**

Appended.

**Supplementary Table 9. Hypertension COX Regression Coefficients**

Appended.

**Supplementary Table 10: GISAID COVID-19 Acknowledgments**

Appended.

